# Cryo-EM Reveals Regulatory Mechanisms Governing Substrate Selection and Activation of Human LONP1

**DOI:** 10.1101/2025.09.05.674599

**Authors:** Jeffrey T. Mindrebo, Lauren Alexandrescu, Jennifer R. Baker, Garret Wang, Gabriel C. Lander

## Abstract

The human AAA+ protease LONP1 plays a central role in maintaining mitochondrial proteostasis. LONP1 processes a vast array of substrates, ranging from damaged or unfolded proteins to specific subunits stably integrated into respiratory complexes. Previous cryo-EM studies of LONP1 uncovered two distinct conformational states corresponding to inactive or active forms of the enzyme. While these states have shed light on the intricacies of LONP1 substrate translocation and proteolytic processing, little is known about the decision-making involved in LONP1 substrate engagement and subsequent initiation of its unfoldase activity. Here, we use cryo-EM to determine a novel ADP-bound, C3-symmetric intermediate state of LONP1 (LONP1^C3^) with putative substrate “fold-sensing” capabilities. Our biochemical and structural data indicate that LONP1^C3^ is an on-pathway intermediate and that is stabilized by interaction with folded substrates. Moreover, we identify additional symmetric and asymmetric conformational states, including a two-fold symmetric split-hexamer conformation, that we associate with the transition from LONP1^C3^ to LONP1^ENZ^. We propose that the C3-state regulates substrate selection and enables LONP1 to efficiently surveil the matrix proteome to ensure selective removal of damaged and dysfunctional proteins as well as privileged LONP1 substrates. These findings collectively provide further mechanistic insights into LONP1 substrate recruitment and engagement and inform on its diverse roles in maintaining homeostasis within the mitochondria.

## Introduction

Mitochondria import 99% of their proteins from nuclear encoded genes using a complex, tightly regulated protein import system that facilitates the translocation of nascent polypeptide chains across one or two lipid bilayers prior to maturation and folding.^1^ Once assembled, these proteins and protein complexes are subjected to high levels of reactive oxygen species produced from oxidative phosphorylation and biosynthetic pathways, such as heme biosynthesis and iron-sulfur cluster biogenesis.^2,3^ Consequently, the mitochondrial matrix is under considerable levels of basal proteotoxic stress. To mitigate this stress, mitochondria possess their own protein quality control network to rectify proteotoxic insults and maintain proteostasis.^4^ A key player in this process is the AAA+ protease LONP1, which is responsible for the removal of oxidized, damaged, and unfolded proteins from the mitochondrial matrix.^5-7^ In addition to proteostasis, LONP1 plays a critical role in diverse mitochondrial processes such as metabolism, genome stability, and responses to hypoxic and oxidative stress.^8-11^ Knockouts of LONP1 are embryonically lethal, and mutations mapped to the LONP1 gene locus have been identified as causal for cerebral, ocular, dental, auricular, skeletal (CODAS) syndrome, a rare multisystem developmental disorder characterized by diverse clinical manifestations.^12^ Due to its central role in stress response, LONP1 is upregulated in most cancers and is implicated in aggressive late-stage malignancies.^13-20^ Therefore, LONP1 is emerging as a master regulator of diverse mitochondrial functions and is of growing interest as therapeutic target.^21-23^

LONP1 assembles as a 600 kDa homohexameric ring from a single polypeptide chain encoding all essential catalytic functions. This assembly consists of an N-terminal substrate binding domain (NTD), a central AAA+ ATPase domain, and a C-terminal protease domain (PD). The NTD can be further subdivided into the NTD globular domain (NTD^GD^), coiled-coil domain (CCD), and a 3-helical bundle (NTD^3H^), which is integrated into the AAA+ domain (**Fig. 1A**).^24^ Recent cryogenic electron microscopy studies (cryo-EM) on bacterial and eukaryotic systems show that Lon proteases adopt two prominent conformations: a proteolytically inactive open conformation or a substrate-bound proteolytically active closed conformation (**Fig. 1B,C**).^25-30^ To adopt the active conformation, the NTD must first engage substrates and facilitate their initial unfolding and transfer to the AAA+ domains (**Fig. 1C**). Once substrate is coordinated by conserved pore loops, the ATPase domains utilize ATP-driven conformational rearrangements to translocate substrate to the PD for subsequent degradation.^26,27,31^

**Figure 1.**
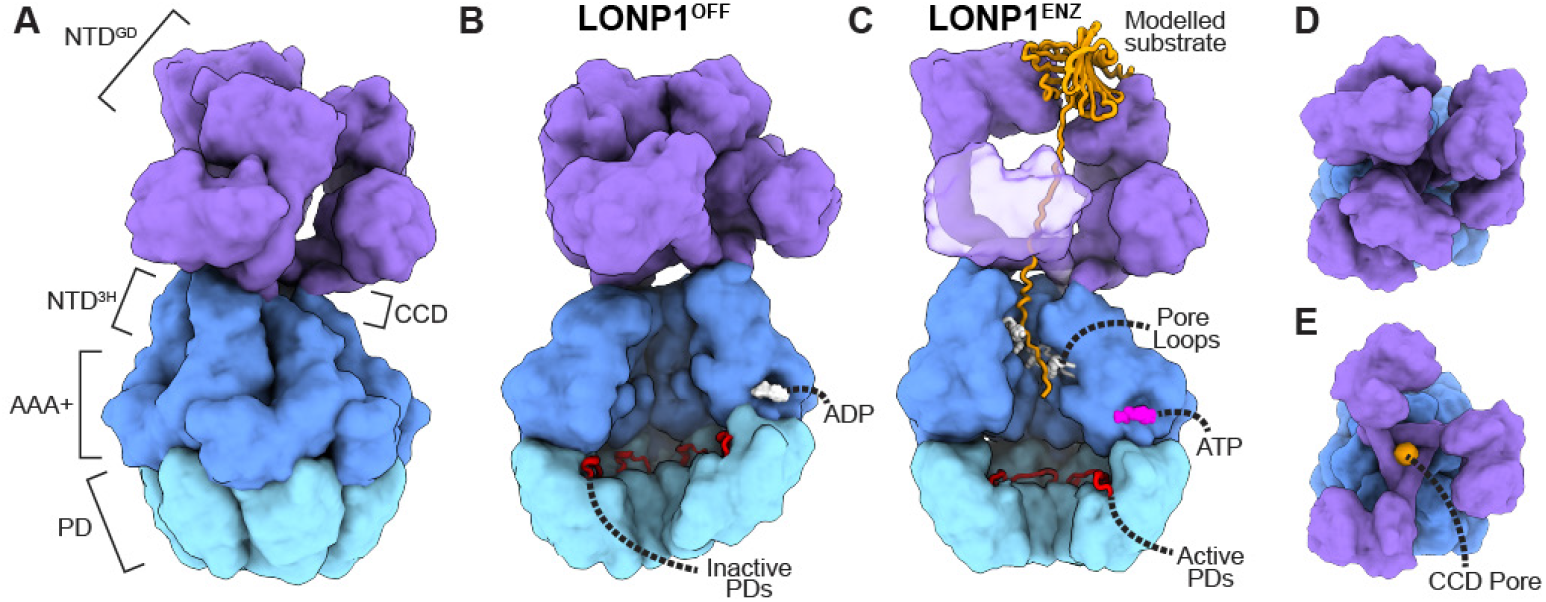
LONP1 structure overview. **A**) Overview of LONP1 structural elements, labeled and colored by domain. The NTD^GD^ and CCD are colored purple, the NTD^3H^ and AAA+ are colored dark blue, and the PD are colored light blue. **B**) The conformation and structural elements that define the inactive open-form of LONP1 (LONP1^OFF^). The ATPase and protease domains adopt a left-handed open lock washer conformation, and all the ATPase domains are bound to ADP (white). The protease domains adopt an asymmetric conformation that positions an inhibitory 3_10_ -helix (red) in the proteolytic active site cleft. **C**) The conformation and structural elements that define the active closed-form of LONP1 (LONP1^ENZ^). Substrate (orange) is bound by the NTD, which then threads through the CCD pore, where it is coordinated by the ATPase pore loops (white). The ATPase domains adopt a right-handed helical conformation that is competent for ATP (magenta) binding and hydrolysis. The PDs adopt a C6-symmetric conformation, resulting in a rearrangement of the inhibitory 3_10_ -helix to form the proteolytic substrate binding site. **D**) Top view of the NTD showing the dimer of trimer arrangement. **E**) The top three NTD domains are removed to show the CCD pore with bound substrate (orange).

Despite numerous recent studies enhancing our understanding of LONP1 structure-function relationships,^26-33^ the processes regulating substrate selection and transition from the open to closed conformation are largely undefined. While it is generally known that the NTD plays a critical role in substrate selection, this domain is conformationally dynamic and difficult to characterize structurally. Numerous low to medium resolution reconstructions of the NTD (**Fig. 1D,E**).^28,31-33^ show how six copies of this domain adopts an interconnected “trimer of dimers” arrangement, which positions three alternating CCD helices to form a triangular motif (CCD pore) above the central axis of the ATPase domains (**Fig. 1D,E**). Substrate-bound structures show continuous polypeptide density traversing this pore and extending to the AAA+ domain where it is coordinated by the conserved pore loops. These observations, along with biochemical analysis, have led to the proposal that the NTD functions as a molecular ratchet coupled to conformational changes in the ATPase domains during substrate translocation.^28,31,32^ Additionally, we recently demonstrated that the NTD also possess additional allosteric binding sites, distinct from the main substrate binding channel, that serve to allosterically regulate function and enhance proteolytic activity.^31^ These findings confirm that the NTD is critical for substrate processing, but do not explain its capacity to regulate substrate recruitment and processing in a complex cellular environment.

In our previous study, we structurally characterized wild type (wt) LONP1 actively processing a model substrate, β- casein, using ATP instead of slowing hydrolysis using ATP analogs or by introducing a Walker B mutation. However, casein has little to no secondary structure and is not a native LONP1 substrate.^31^ To better inform on the underlying regulatory mechanisms employed by LONP1 to selectively degrade structured, or partially structured substrates, we used cryo-EM to visualize LONP1 degrading bona fide human matrix localized protein substrates: steroid acute regulatory protein (StAR) and mitochondrial transcription factor A (TFAM).^34,35^ These experiments led to the discovery of a novel ADP-bound C3-symmetric conformation of LONP1, LONP1^C3^, that integrates structural motifs from the NTD and ATPase domains to regulate substrate access to the AAA+ motor. We demonstrate that the C3-symmetric conformation is an on-pathway intermediate critical for function and that it is preferentially stabilized by substrates with higher-order structure. Additionally, we identify a C2-symmetric split-hexamer conformation and an asymmetric right-handed spiral conformation that represent plausible conformational intermediates spanning the transition from LONP1^C3^ to the active substrate-bound conformation, LONP1^ENZ^. Importantly, LONP1^C3^ provides a mechanism to regulate substrate selection and processing at early stages of recruitment prior to peptide engagement and unfolding by the ATPases. We propose that the C3-state operates as checkpoint to ensure the selective removal of damaged/unfolded and privileged LONP1 substrates from the mitochondrial matrix. The findings in this study provide much-needed insight into LONP1 regulation and provide the groundwork to inform LONP1’s role in other critical mitochondrial processes.

## Results

### A Novel C3-Symmetric ADP-bound Regulatory State of LONP1

In our previous study, we observed LONP1 actively degrading a model substrate, β-casein, in the presence of ATP.^31^ However, casein is not a native substrate of LONP1 and has little to no secondary structure. Therefore, we sought to visualize early stages of substrate recruitment and selection by evaluating LONP1 degrading StAR, a verified, folded LONP1 substrate.^34,36^ To visualize LONP1 actively processing substrate, we preincubated LONP1 and StAR at equimolar ratios and rapidly plunge froze samples after adding ATP. Image analysis of our LONP1+StAR dataset showed the presence of the open- and closed-forms of LONP1, although 3D-classification of the closed-form yielded two classes with distinct conformational states (**Fig. S1**). The first class, resolved to ∼ 3.3 Åresolution, is nearly identical to our previous LONP1^ENZ^ structures (PDB ID) representing the fully activated LONP1 holoenzyme (**Fig. S1**). Further refinement of the second class surprisingly yielded a previously unobserved conformation of LONP1 bearing C3-symmetry (LONP1^C3^). The overall conformation of this symmetric state is distinct from both LONP1^ENZ^ and LONP1^OFF^, with a better resolved CCD subdomain of the NTD positioned above the pore (**Fig. 2A**). Refinements using C1-symmetry resulted in a ∼3.2 reconstruction while application of C3-symmetry increased the nominal resolution to ∼2.9 Å.

**Figure 2.**
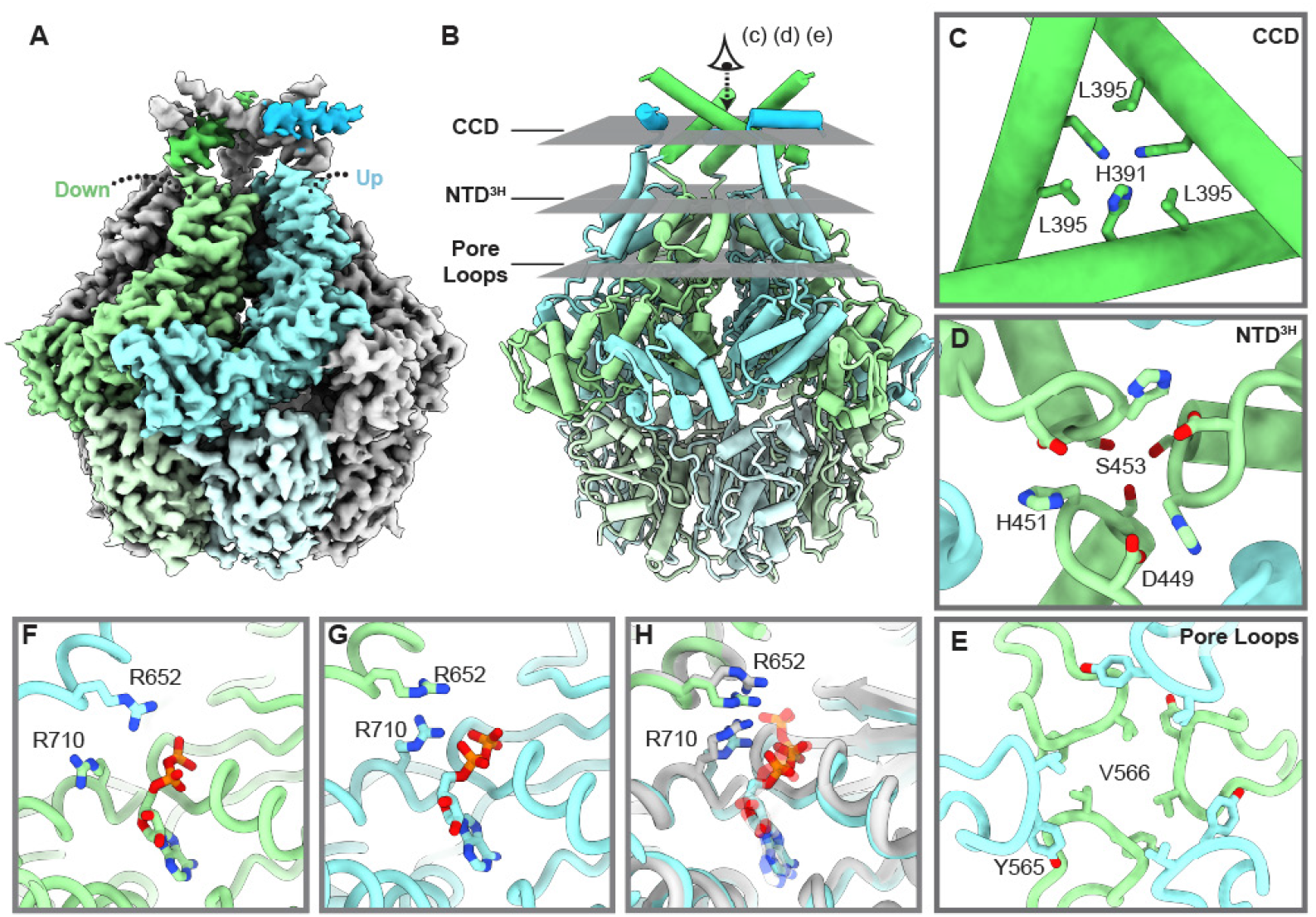
Structural overview of LONP1^C3^ state. **A**) Density for the LONP1^C3^ map is shown with the asymmetric unit colored in shades of blue (down conformation) or green (up conformation) by domain. The color shades denote domain architecture (light is PD, medium is AAA+ and NTD^3H^, and dark is CCD). **B**) Overview of the LONP1^C3^ atomic model with key areas of regulation highlighted by the grey planes. The perspective eye provides the viewing angle for panels C-E. **C**) Conformational state of the CCD subdomain in LONP1^C3^. Residues H391 and L395 form a tightly packed core that prohibits entrance through the pore. **D**) The conformational state of the NTD^3H^ subdomains of the three downward positioned subunits. **E**) Alternating up (green) and down (blue) pore loop conformations of the ATPase domains. **F**) The down-conformation ATPase active site bound to ADP. **G**) The up-conformation ATPase active site bound to ADP. **H**) Structural overlay of the up-conformation ATPase subunit (light green) with a LONP1^ENZ^ ATP-bound subunit (white) demonstrating the similarity in active site conformation between the two nucleotide states.

Similar to other observed conformations of LONP1, the NTD^GD^ subdomains of LONP1^C3^ were poorly resolved. However, the rest of the structure, including the triangular CCD of the NTD positioned above the central pore, was sufficiently resolved for atomic modeling. While LONP1^C3^ exhibited the same catalytically active C6-symmetric PD conformation as LONP1^ENZ^ (**Fig. S2**), the ATPase domains of LONP1^C3^ adopt a conformation that differs from all previously characterized LONP1 conformers. The ATPases are a defining feature of the three-fold symmetric state, adopting an alternating up-and-down arrangement around the ring, with three subunits tilted up and away from the central axis and the other three gathered together at the central pore (**Fig. 2A,B**). Notably, Åwe were able to confidently discern the helical register and position relevant side chains of the CCD (**Fig. 2A,B**), which is in a more tightly compact conformation than our previous substrate-bound reconstructions.

Whereas the CCD of LONP1^ENZ^ positions a conserved tyrosine residue at the central pore for interaction with a bound substrate, the triangular CCD of LONP1^C3^ is arranged to prohibit transfer of substrates from the NTD to the AAA+ motor. Viewed along the symmetry axis from above, residues His391 and Leu395 from each central CCD helix tightly pack together and sterically block incoming substrate from accessing the ATPase channel (**Fig. 2C, S3**). Directly below the CCD, the NTD^3H^ helical bundles of the down-subunits are positioned in proximity of one another to form an electrostatic interaction network at the central axis that further limits access to the AAA+ channel (**Fig. 2D**). The conserved pore loops of the AAA+ domains within the channel are intercalated in an alternating up-and-down configuration, creating a rigid network unable to coordinate with substrate (**Fig. 2E**). Finally, all six ATPase active sites are bound to ADP, despite the up-subunits being in a conformation that is consistent with previously observed ATP-bound subunits (**Figs. 2F-H, S4**). This observation that the nucleotide binding pocket of the up-conformation is nearly identical to the ATP-bound form suggests that ADP-bound subunits can adopt diverse conformations to facilitate function or regulation (**Fig. 2H**). The series of vertically layered structural elements, in concert with ADP-bound AAA+ domains, present an assembly that is incompatible with substrate engagement and translocation. Given that LONP1 must make molecular decisions to degrade, chaperone, or release mitochondrial matrix proteins, we posit that the LONP1^C3^ conformation serves as a checkpoint or fold-sensing state by blocking access to the AAA+ and PDs, prohibiting processing of off-target substrates.

### The C3-state is important for function

To evaluate the relevance of the LONP1^C3^ conformation in enzymatic function, we prepared a panel of mutations that we expect to perturb interactions stabilizing the CCD and NTD3H in the C3-state. Mutants were subjected to both an NADH-coupled ATPase assay and a FITC-casein degradation assay. Importantly, LONP1 exhibits both basal and stimulated ATPase rates in the absence or presence of added substrate, respectively. We surmise that the basal rate represents LONP1’s ability to degrade partially unfolded LONP1 particles or contaminants present in the in the sample. Addition of a suitable substrate leads to higher levels of engagement, resulting in an increase in the total population of particles actively consuming ATP and degrading substrate.

We first mutated both CCD residues, His391 and Leu395, to alanine, and found that the double mutant had little effect on proteolytic degradation and ATP consumption (**Fig. 3**). However, mutating these residues to either a charged residue (glutamate) or a bulky residue (tryptophan), which would likely hinder adoption of the C3-conformation, substantially altered LONP1 activity. The H391E mutation nearly abolishes activity, and H391W shows a ∼ 15-fold reduction in both ATPase and FITC-casein degradation activity (**Fig. 3**). These results suggest that transitions through the LONP1^C3^ conformation are important for substrate processing.

**Figure 3.**
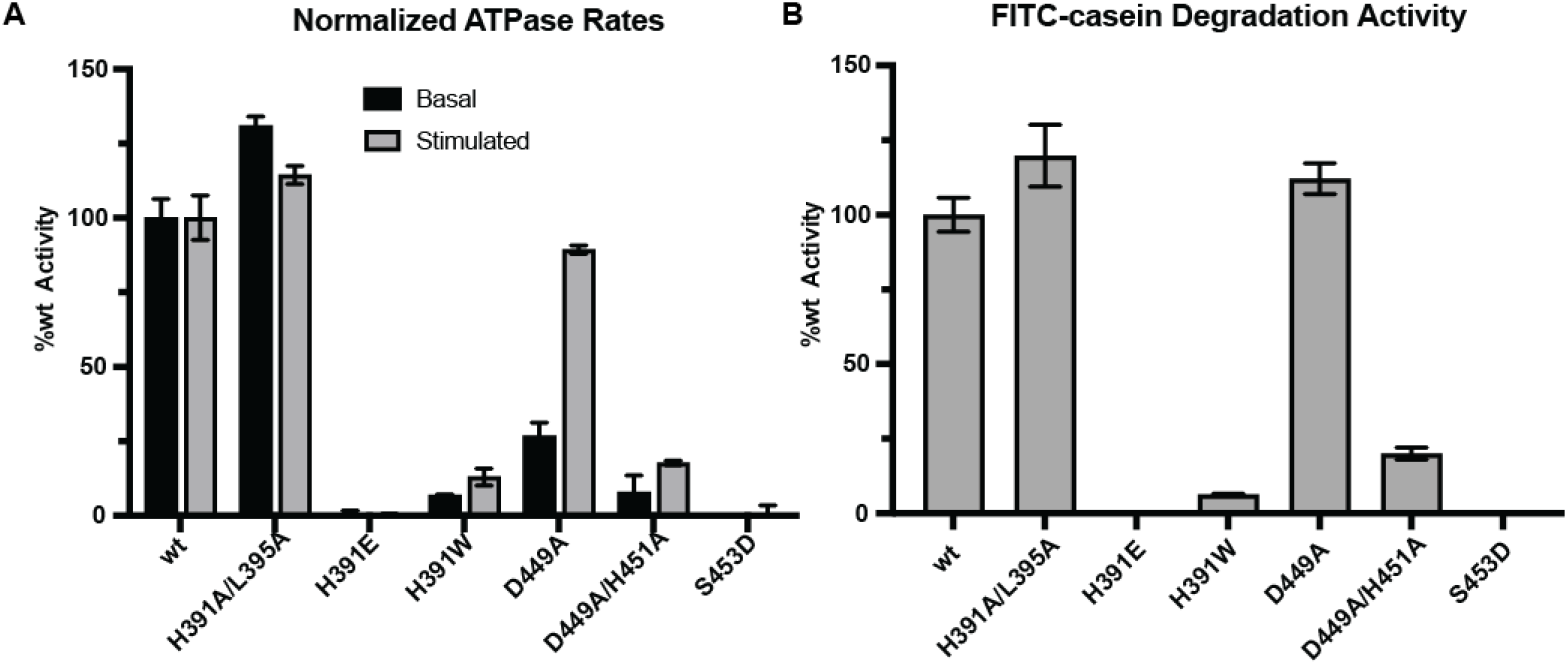
ATPase and FITC-casein activity assays. **A**) Basal and stimulated (with 10 µM casein) ATPase rates of wt and mutants normalized to wt activity. **B**) FITC-casein degradation rates of wt LONP1 and mutants normalized to wt activity. All assays were repeated in biological triplicate and error bars represent standard deviation of the mean.

We next characterized interactions at the NTD^3H^. In all prior cryo-EM structures of nucleotide-bound LONP1, Asp449, His451 and Ser453 are not observed to interact with substrate or nucleotide. Further, unlike the CCD, this region has no prior implications in substrate recruitment or processing. Thus, we expected that mutating these residues would inform on their specific role in adopting the C3-state. A single point mutant, D449A, showed nearly wt activity for stimulated ATPase and FITC-casein degradation rates (**Fig. 3**), but the D449A/H451A mutant showed a ∼5-fold reduction in FITC-casein degradation activity and ∼6-fold reduction in stimulated ATPase rate (**Fig. 3**). To more intensely disfavor the C3-state, we introduced a S453D mutation, which would increase the bulkiness of this position and lead to an overabundance of negative charge, thereby preventing the three NTD3H domains from coming together at the central channel. Accordingly, we observed the S453D mutant to be inactive in our assay conditions for both FITC-casein and ATPase activity (**Fig. 3**). Together, these results demonstrate that perturbing the LONP1^C3^ conformation negatively impacts function.

### LONP1^C3^ represents an on-pathway intermediate

Structurally, the C3-state blocks substrates from accessing the protease sites through the ATPase domains, and could therefore serve as an important regulatory step in holoenzyme activation. Our assay results suggest that adopting the C3-state plays a role in proteolytic and ATPase activity, although it is unclear if the C3-state is an on-pathway intermediate adopted as LONP1 progresses from the inactive open form to the active closed form. We know that LONP1 exists as an ensemble of conformations where the open, closed, and now C3-state are present during substrate processing. Therefore, the ATPase and proteolytic degradation rate are likely determined by the percentage of LONP1 particles in the active closed form, as the open-form and C3-state are unable to participate in ATP hydrolysis or substrate translocation. If the C3-state serves as a checkpoint during substrate processing, disrupting transition through the C3-state should manifest as a lower population of actively translocating LONP1 particles. In this case, ATPase kinetic parameters would have a lower k_cat_ but little or no change in the K_M_.

To evaluate this hypothesis, we determined the catalytic parameters of stimulated ATPase activity for wt LONP1 and the D449A/H451A mutant. Intriguingly, the D449A/H451A mutant had a 4.7-fold reduction in k_cat_ while the K_M_ slightly decreased (**Fig. 4A**), suggesting that disrupting the LONP1^C3^ conformer hinders adoption of the proteolytically competent closed-form, thereby decreasing LONP1’s ability to hydrolyze ATP. This conclusion is further supported by the nearly equivalent drop in FITC-casein degradation rate and ATPase rate for the D449A/H451A mutant, suggesting these activities are coupled (**Fig. 3**). Therefore, we propose that destabilizing the LONP1^C3^ conformation disrupts LONP1’s capacity to adopt the active form and process proteolytic substrates.

**Figure 4.**
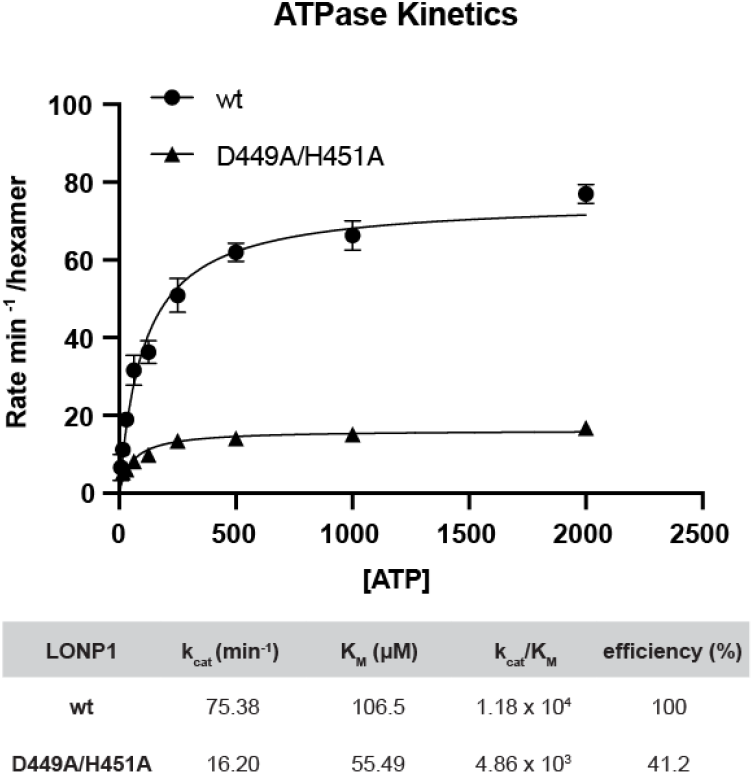
ATPase kinetics wt and D449A/H451A double mutant. Michaelis-Menten Kinetics of wt LONP1 and D449A/H451A ATP-ase activity. Table below graphical data provides catalytic parameters for wt and D449A/H451A double mutant.

### The CCD conformation of LONP1^OFF^ resembles LONP1^3^

We next sought to determine the structural arrangement of the CCD in the LONP1^OFF^ conformation to determine its relationship to LONP1^C3^ and LONP1^ENZ^. We identified a subset of particles within a large stack of open-conformation particles in our LONP1+StAR dataset bearing improved density for the sixth subunit and CCD domain (**Fig. S3**). Although the resolution of this region was lower than observed in the LONP1^C3^ and LONP1^ENZ^ reconstructions, it was sufficient to rigid-body fit atomic models into the density for a structural comparison of the three states. The CCD adopts a notably tighter conformation in LONP1^OFF^ than in LONP1^ENZ^, more closely matching that of the LONP1^C3^ conformation, suggesting that in the absence of substrate the CCD conformation is consistent with a narrowed CCD pore (**Fig. S4**). Based on their structural similarity, it is reasonable that the NTD of LONP1^OFF^ could transition to the C3-state as an on-pathway intermediate as LONP1 progresses to the substrate-engaged closed conformation.

### Stabilization of LONP1^C3^ is substrate-dependent

Our data suggest that the LONP1^C3^ state represents a functionally relevant point on the conformational landscape associated with holoenzyme activity. We speculate that this symmetric organization could serve as a checkpoint involved in determining the fate of an encountered substrate. In our previous studies using β-casein, we did not observe the LONP1^C3^ conformer, suggesting that the substrate “foldedness” could be a factor in stabilizing this conformation. To determine if the formation of LONP1^C3^ is substrate dependent, we performed an additional cryo-EM experiment using the mitochondrial transcription factor A (TFAM) instead of StAR. TFAM is a known *in vivo* LONP1 substrate and fits a molten globule model with predominantly helical secondary structure in the absence of DNA,^35^ serving as a model substrate exhibiting an intermediate degree of foldedness as compared to β-casein (unstructured) or StAR (structured) (**Fig. S5**).

Analysis of the TFAM+LONP1 dataset revealed a mixture of LONP1^OFF^, LONP1^ENZ^, and LONP1^C3^ states (**Fig. S6A**), suggesting that LONP1 processing of intermediate- or well-folded substrates leads the enzyme to adopt a C3-state (**Fig. 5A**). In addition to the active form and C3-state, we also identified two additional ADP-bound states that resemble LONP1^C3^ but do not fully adhere to 3-fold symmetry. These asymmetric intermediates have reduced density for the CCD and greater heterogeneity in the NTD^3H^ subdomain and pore loops (**Figs. S7, S8A**) (discussed further in the following section).

**Figure 5.**
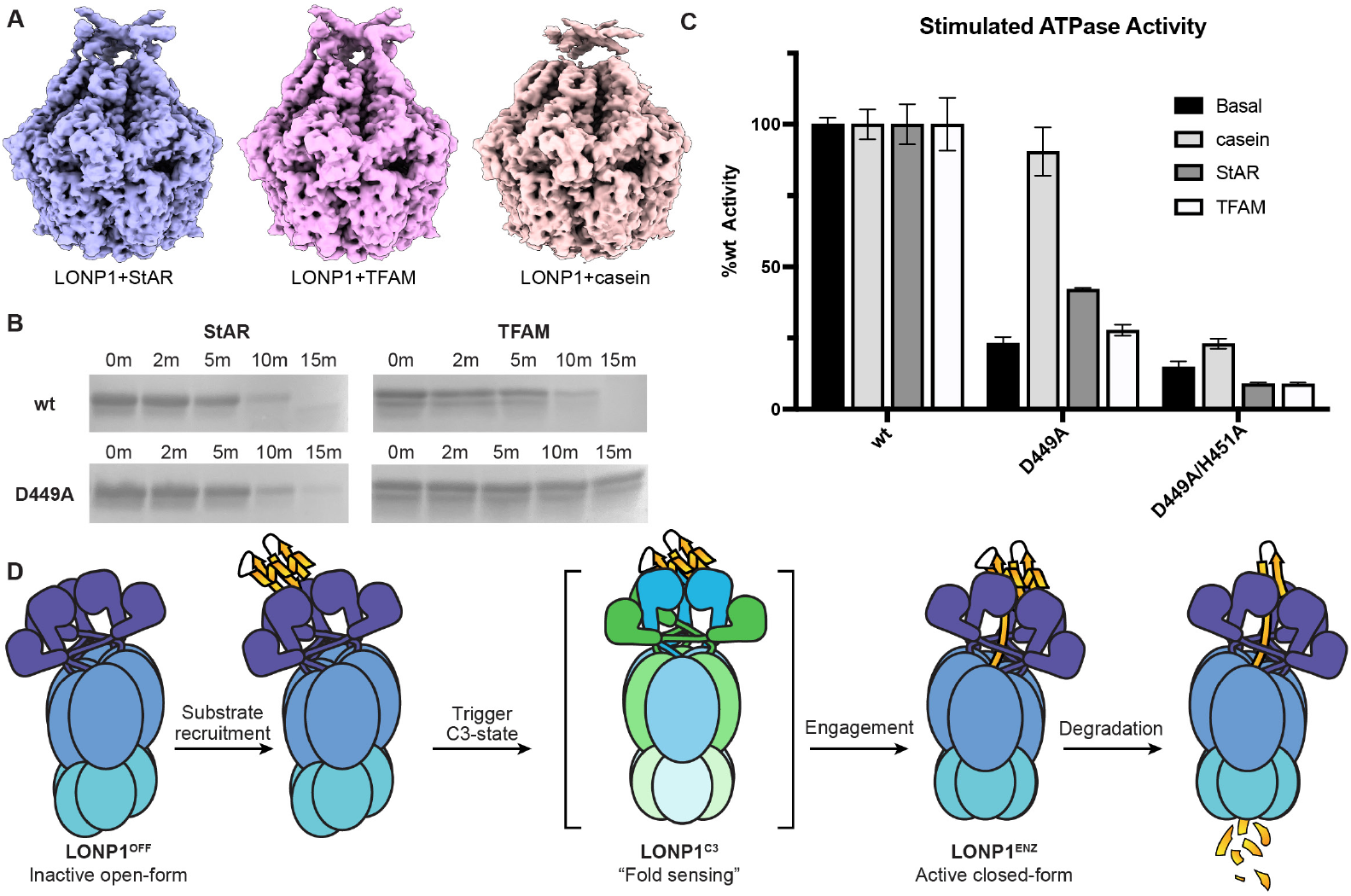
Substrate dependence of C3-state. **A**) Comparison of LONP1^C3^ conformational states across datasets of LONP1 degrading either StAR, TFAM, or β-casein. The LONP1-casein structure was obtained after applying symmetry expansion followed by 3D-classifiction, whereas LONP1-StAR and LONP1-TFAM were readily identified using standard processing pipelines (**Figs S1, S6**). Binarization threshold was set to 0.13 for all volumes. **B**) Gel-based degradation assay of StAR and TFAM with either wt LONP1 or the D449A mutant. The D449A mutant presents reduced degradation of StAR and TFAM despite wt activity levels against unstructured β-casein. Gel-based assays were repeated in duplicate. **C**) Substrate-dependent ATPase stimulate rates of wt LONP1, D449A, and D449A/H451A. Results indicate that the relative-to-wt ATPase stimulation is substrate dependent with mutations that destabilize LONP1^C3^. ATPase stimulation assays were repeated in biological triplicate and error bars represent standard deviation of the mean. **D**) Proposed mechanism where LONP1^C3^ is an on-pathway intermediate between the LONP1^OFF^ and LONP1^ENZ^.

In our previous study where LONP1 was incubated with the unstructured model substrate β-casein, we did not identify the C3-state in our cryo-EM dataset. However, reprocessing this data using the workflow we established for LONP1+TFAM revealed a subset of particles consistent with the LONP1^C3^ conformation, which we resolved to ∼ 3.2 Åresolution (**Fig. S6B**). While the overall structure of this β-casein-associated LONP1^C3^ was consistent with those observed in the presence of StAR or TFAM, the CCD and NTD^3H^ sub-domains are not as well ordered (**Fig. 5A**). This finding is consistent with LONP1 adopting the C3-state during substrate processing, and that the stability of the CCD domain correlates with the foldedness of the substrate. Therefore, transitions through the C3-state may occur rapidly with unstructured or unfolded substrates but persist with more folded substrates such as TFAM and StAR. Thus, we posit that the CCD serves as a structural checkpoint, giving LONP1 the ability to sense substrate foldedness and specifically target proteins that present suitable degrons for proteolysis.

To support this hypothesis, we evaluated the degradation of folded substrates by wt LONP1 and our D449A mutant, which we expect to disrupt the C3 state, but observed to maintain wt levels of casein-stimulated ATPase activity and FITC-casein degradation activity (**Fig. 3**). wt LONP1 degrades StAR or TFAM to completion in under 15 minutes in a time-course degradation assay (**Fig. 5B**). The D449A mutant, however, showed a notable reduction in degradation activity for both StAR and TFAM (**Fig. 5B**). We also observed the same substrate-dependent trend in ATPase activity for the D449A and D449A/H451A mutants, where the mutations that putatively disrupt the C3-state showed reduced ATP hydrolysis for StAR and TFAM relative to casein (**Fig. 5C**). These findings collectively support a model where the LONP1^C3^ conformation operates as a checkpoint to regulate substrate selection and processing (**Fig. 5D**), and that transitioning through this C3-symmetric conformation aids in processing folded substrates.

### A C2-symmetric split-hexamer intermediate bridges LONP1^C3^ and LONP1^ENZ^ states

We were intrigued by the presence of C3-like intermediate states in our datasets, which prompted us to further process these images in search of additional LONP1 conformations. Further analysis of the particles contributing to the intermediate conformations identified in the LONP1+TFAM dataset revealed the presence of a LONP1 conformation bearing two-fold symmetry (**Figs. 6A,B, S7, S8B**). We refined the subset of particles associated with this conformation with imposed C2-symmetry, resulting in a ∼ 3.4 Åresolution reconstruction that we refer to as LONP1^C2^ (**Figs. S7, S8B**).

**Figure 6.**
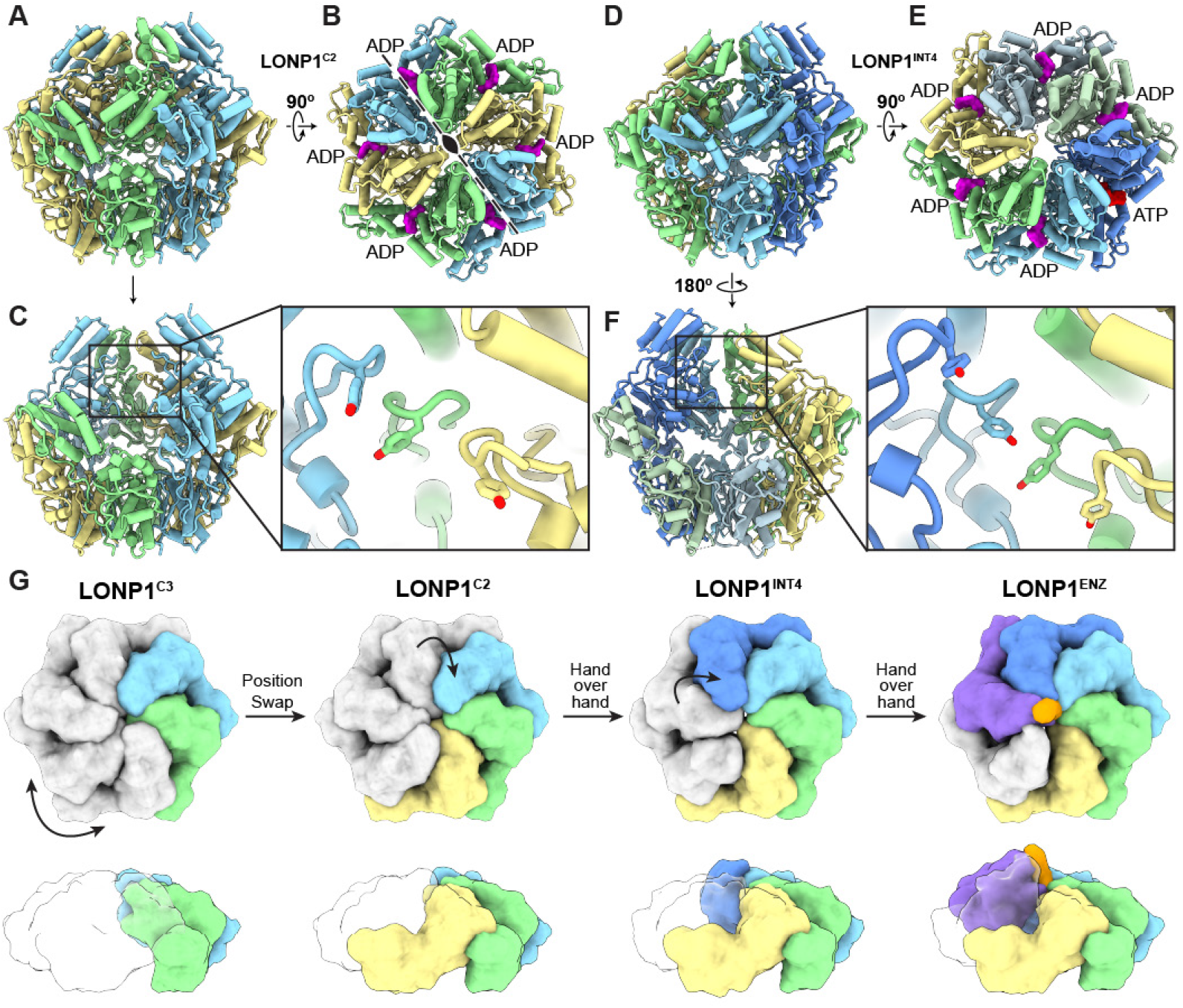
Structural overview of LONP1 processing intermediates. **A**) Side view of LONP1^C2^ emphasizing the three-subunit right-handed ATPase spiral, with yellow at the bottom and light blue at the top. **B**) Top view of LONP1^C2^ demonstrating the C2-symmetry axis and nucleotide state of each ATPase domain. **C**) Cutaway and close-up view of the pore loops of the rear three-subunit right-handed spiral. **D**) Side-view of LONP1^INT4^ showing the four-subunit right-handed spiral. **E**) Top view of LONP1^INT4^ showing the more open central pore relative to LONP1^C2^ and nucleotide state of each ATPase domain. **F**) Cutaway and close-up view of the pore loops of the rear ATPase domains in LONP1^INT4^ forming a four-subunit right-handed spiral. **G**) Proposed order of conformational arrangements to transition from LONP1^C3^ to LONP1^ENZ^.

In the LONP1^C2^ conformation, the fully ADP-bound ATPase hexamer is split into two symmetric half-spirals (**Figs. 6A-C** and **S8B,F**), each of which is superimposable with the three lowermost ATPase domains that form the right-handed spiral in LONP1^ENZ^ (**Fig. S8C**). This organizational similarity is evident when comparing the pore loop residues of the LONP1^C2^ and LONP1^ENZ^ structures (**Fig. S8C**). The presence of a right-handed staircase organization in the LONP1^C2^ conformation is somewhat surprising, as this structure lacks any appreciable bound substrate density in the central pore (**Fig. S8B,D**), yet substrate has been proposed to play a role in organizing the right-handed staircase organization of ATPase domains in AAA+ proteases.^26,31^ The LONP1^C2^ conformer also bears a structural consistency with the LONP1^C3^ conformation, as the subunits at the interface of each half-spiral maintains the “up-down” configuration of the neighboring ATPase domains observed in the LONP1^C3^ conformation. We note that the only structural difference between the LONP1^C3^ and LONP1^C2^ conformations is that two of the protomers at one of the half-spiral interfaces have flipped their orientation from down-up to an up-down arrangement (**Fig. S8E, Movie 1**). Thus, by undergoing a relatively minor conformational rearrangement of two subunits, LONP1 can switch from the C3-to the C2-symmetric arrangement. In the context of the emerging LONP1 conformational landscape, the LONP1^C2^ conformer represents an intermediate conformation that bridges the transition from LONP1^C3^ to LONP1^ENZ^. Such sequential symmetric rearrangements would allow redundancy in substrate engagement through interaction with symmetrically equivalent subunits, enhancing efficiency of the activation process.

Further analysis identified two asymmetric intermediate states whose ATPases are arranged in sequential conformational steps representing the transition from the LONP1^C2^ split-hexamer state to the LONP1^ENZ^ full right-handed spiral (**Fig. S9**). For LONP1^INT3^, we observe a notable break in the symmetry where the ATPase domain that is counter-clockwise adjacent to the three-subunit right-handed spiral adopts a higher position (dark blue in **Fig. 6G**), beginning the formation of a four-subunit right-handed spiral (**Fig. S9**). LONP1^INT4^ continues this process by fully establishing the four-subunit spiral, where the subunits are positioned as they would be in the context of the canonical five-subunit AAA+ spiral (**Figs 6D-G, S9**). Notably, this integration of the fourth subunit into the LONP1^INT4^ spiral is consistent with the movement of this subunit during the canonical hand-over-hand conformational rearrangements thought to be used by AAA+ domains to facilitate substrate translocation. However, despite adopting a staircase arrangement, the pore loops are not engaged with substrate in either of these structures (**Fig. 6F, S10**). Further, all subunits contain nucleotide densities consistent with ADP, with the exception of the uppermost subunit of LONP1^INT4^, which contains ATP (**Figs. 6B, S11**). This ATP occupancy may indicate that the step-wise activation from LONP1^C2^ to LONP1^ENZ^ is encouraged by nucleotide exchange to fully stabilize the right-handed spiral for substrate engagement. Finally, the LONP1^INT4^ conformation can readily transition to the canonical right-handed spiral through an additional subunit rearrangement to position a fifth protomer atop the right-handed spiral (purple in **Fig. 6G**), completing formation of the LONP1^ENZ^ conformation compatible with substrate translocation (**Movie 1**). Based on these structural analyses, we propose that the activation mechanism of LONP1 begins with substrate engagement at the NTD’s in the lefthanded asymmetric LONP1^OFF^ state, which then progresses through a series of symmetric and then asymmetric rearrangements to ultimately establish the right-handed asymmetric substrate-bound active-form, LONP1^ENZ^ (**Fig. 6G, Movie 1**).

## Discussion

A critical aspect of mitochondrial protein quality control is the selective degradation of damaged, unfolded, or aggregated proteins, which if not removed, perturb mitochondrial fitness and function. To remove damaged proteins, mitochondria harbor four distinctly targeted AAA+ proteases. Two of these AAA+ proteases, AFGL32 and YME1L, are membrane-bound and face either the matrix or the intermembrane space, respectively, while LONP1 and CLPXP are soluble matrix localized proteins.^3^ Both LONP1 and CLPXP are responsible for the removal of damaged and misfolded matrix localized proteins, but knockouts of CLPP are tolerated while LONP1 knockouts are embryonically lethal.^12,37-40^ Moreover, in *S. cerevisiae*, ClpP is absent from the genome, leaving LONP1 as the sole AAA+ protease surveilling the mitochondrial matrix.^39^ Therefore, CLPXP, while important for metabolism and stress response, appears to play a secondary role in maintaining proteostasis in the matrix, whereas LONP1 is required.^37,40,41^ This raises an important question: How is LONP1 able to selectively remove the bulk of damaged and unfolded proteins while also recognizing privileged folded substrates, such as TFAM and StAR?

To address this question, we used cryo-EM to characterize LONP1 actively degrading folded substrates at physiologically relevant temperatures. With this approach, we identified and biochemically characterized a novel C3-symmetric state of LONP1 that integrates structural components from the NTD and AAA+ domains to regulate substrate engagement by the ATPase. Importantly, this mode of regulation is different than the previously reported LONP1^OFF^ state, where the proteolytic domains adopt an inactive conformation to limit off-target degradation when LONP1 is not engaged with substrates. Therefore, LONP1^C3^ regulates function at a more critical stage in the decision-making process by preventing translocation rather than proteolysis. We propose that a core function of LONP1^C3^ is to serve as checkpoint to determine whether to degrade or release a bound substrate. Such a checkpoint would allow LONP1 to evaluate the state of matrix proteins and only use ATP to unfold damaged proteins or privileged substrates. In support of this notion, we demonstrate that the adoption of LONP1^C3^ is critical for substrate processing, and that it is preferentially stabilized in the presence of folded substrates. Moreover, our D449A mutant presents wt levels of proteolytic activity with casein but has reduced activity against more challenging substrates like TFAM and StAR. Taken together, these findings indicate that less structured substrates, such as unfolded matrix proteins, would rapidly transition through the C3-state, while more structured substrates persist in the C3-state until a suitable degron is engaged, or the protein is instead released.

In addition to identifying and biochemically evaluating LONP1^C3^, we also elucidated additional intermediate states that are present during the active degradation of all three tested substrates. Specifically, our LONP1^C2^ conformation, along with LONP1^INT3^ and LONP1^INT4^, provide a full perspective on the transitions required to adopt the canonical active closed-form, LONP1^ENZ^. These data lead us to propose a reaction scheme where LONP1^C3^, and the additional intermediate states, are on-pathway between LONP1^OFF^ and LONP1^ENZ^. We propose a mechanism where substrate binding to the NTD of LONP1^OFF^ triggers the adoption of the LONP1^C3^ conformation, which performs the initial unfolding events to facilitate degron engagement. If the NTD is unable to engage a suitable degron on the bound substrate, it dissociates and LONP1^C3^ transitions back to LONP1^OFF^. If the bound substrate is to be degraded, there is a progressive relaxation of symmetry from LONP1^C3^ to LONP1^C2^, where each asymmetric unit is composed of a three-subunit right-handed spiral. At this stage, a series of hand-over-hand-like rearrangements^26,27,31^ establish the asymmetric right-handed spiral, transitioning through LONP1^INT3^ and LONP1^INT4^, to finally arrive at LONP1^ENZ^. In this way, LONP1 can transition from an asymmetric, inactive left-handed conformation to an asymmetric, active right-handed conformation through a series of symmetric intermediates. Importantly, these transitions are seemingly mediated by the largely unresolved NTD, which similarly to other structures, is only visible at low-resolution for LONP1^C3^.

While we were unable explicitly identify bound substrate in our activation intermediates, these transitions are likely coupled to the substrate recognition and engagement process, as demonstrated by our biochemical assays. Therefore, the question remains as to how these rearrangements are coordinated across the assembly. Interestingly, the NTD^3H^, which is unique to LONP1, appears to function as a signal transducer to facilitate allosteric communication from the NTD to the ATPase domains below. The NTD^3H^ is integrated into the ATPase cassette by a significant hydrophobic interface, which we propose couples the movement of the ATPase domains with rearrangements occurring in CCD domain above (**Fig. S12**). LONP1 thus employs a sophisticated allosteric mechanism to select for specific substrates in a complex cellular environment in an ATP-independent manner, simultaneously optimizing energy consumption and enforcing substrate selectivity. Additionally, these newly reported conformational intermediates, particularly the LONP1^C3^ conformation, could serve additional roles in other reported LONP1 activities, such as DNA binding or chaperone activity.^42-45^ In such cases, regulating access to the AAA+ motor could be critical to prevent unintended off-target degradation of client proteins or DNA-bound protein assemblies. Therefore, we are hopeful that the discovery and validation of a novel regulatory state, and our proposed activation mechanism, will provide avenues for future studies to determine LONP1’s role in regulating diverse aspects of mitochondrial biology in human health and disease.

## Materials and Methods

### Construct generation

TFAM and StAR were synthesized without their mitochondrial targeting sequences by Twist Biosciences and cloned into pET-28a expression vector using NCO1 and XHO1 cut sites. All mutations were generated using the method described by Liu and Naismith.^46^

### Protein expression

LONP1 was transformed into Rosetta (DE3) PlysS cells (Sigma). 20 ml cultures were grown overnight and used to inoculate 1 L of terrific broth media (VWR). Cells were grown until they reached an OD of 0.8 and then cooled on ice before transferring to a 16 ^*°*^C incubator for overnight induction using 0.5 M IPTG (Sigma). Cells were harvested by centrifugation and frozen until purification. To purify LONP1, cell pellets were resuspended in lysis buffer (50 mM Tris pH 8.0, 300 mM NaCl, 10% glycerol) to a volume of 40 ml. Cells were lysed by sonication and lysate was clarified at 30,000 X G for 45 minutes. Lysate was batch bound to nickel resin (Qiagen) for 30 minutes before applying to a column. The nickel beads were washed with 50 ml of lysis buffer, 50 ml of lysis buffer supplemented with 20 mM imidazole, and 30 ml of lysis buffer supplemented with 50 mM imidazole. Proteins were eluted off the column in lysis buffer supplemented with 250 mM imidazole. Fractions containing protein were pooled, concentrated, and injected onto a superose 6 increase 10/300 GL FPLC column (Cytiva) equilibrated with protein storage buffer (50 mM Tris pH 8.0, 150 mM NaCl, 0.5 mM TCEP). Fractions containing hexameric protein were pooled, concentrated, and flash frozen for future use. The same protocol was used for the expression and purification of TFAM and StAR except that they were further purified using a superdex S200 10/300 GL FPLC column (Cytiva).

LONP1 was expressed in *E. coli* Rosetta (DE3) pLysS cells (Sigma) following transformation. Starter cultures (20 ml) were grown overnight and subsequently used to seed 1-liter volumes of terrific broth (VWR). Cultures were incubated until reaching an optical density of 0.8, then chilled on ice and shifted to a 16 ^*°*^C incubator for overnight induction with 0.5 M IPTG (Sigma). After induction, cells were harvested by centrifugation and stored frozen prior to purification. For purification, frozen cell pellets were thawed and resuspended in 40 ml of lysis buffer containing 50 mM Tris pH 8.0, 300 mM NaCl, and 10% glycerol. Cell disruption was achieved by sonication, and the lysate was clarified via centrifugation at 30,000 × g for 45 minutes. The cleared lysate was batch incubated with nickel affinity resin (Qiagen) for 30 minutes, then transferred to a column format. Resin was sequentially washed with 50 ml of lysis buffer and lysis buffer with 20 mM imidazole followed by one additional wash of 30 ml of lysis buffer containing 50 mM imidazole. Protein elution was performed using lysis buffer supplemented with 250 mM imidazole. Eluted fractions containing LONP1 were pooled and concentrated, then subjected to size-exclusion chromatography on a Superose 6 Increase 10/300 GL column (Cytiva) pre-equilibrated with storage buffer (50 mM Tris pH 8.0, 150 mM NaCl, 0.5 mM TCEP). Hexameric fractions were combined, concentrated, and snap-frozen for later use. TFAM and StAR proteins were expressed and purified following the same protocol, with the exception that size-exclusion purification was carried out using a Superdex 200 10/300 GL column (Cytiva).

### Coupled ATPase assay

ATPase activity was assessed using an enzyme-coupled assay that links ATP hydrolysis to NADH oxidation via lactate dehydrogenase (LDH). In this setup, pyruvate kinase (PK) catalyzes the regeneration of ATP from ADP by converting phosphoenolpyruvate (PEP) to pyruvate. The resulting pyruvate is then reduced to lactate by LDH, consuming NADH in the process. The depletion of NADH was tracked by monitoring fluorescence loss (excitation at 340 nm, emission at 465 nm), which corresponds to its conversion to NAD^+^. Assay reactions began with the preparation of a 2X mastermix containing 2.0 mM PEP (1PlusChem), 0.4 mM NADH (Cayman Chemical), 60 U/ml LDH (Worthington Chemical), and 20 U/ml PK (Sigma). For kinetic analysis, the mastermix was diluted to 1X in reaction buffer consisting of 100 mM KCl, 50 mM Tris pH 8.0, and 10 mM MgCl_2_, followed by a 10-minute pre-incubation at 37 ^*°*^C with the desired ATP concentration. Reactions were initiated by adding enzyme to a final concentration of 42 nM for wild-type LONP1, H391A/L395A, D449A, D449A/H451A, 166 nM for H391W, and 333 nM for H391E and S453D. Fluorescence data were analyzed using linear regression to obtain reaction rates, which were converted to molar units based on a standard curve generated from known NADH concentrations. Background rates—measured in the absence of enzyme—were subtracted from the raw rates to yield net ATPase activity. For Michaelis-Menten kinetics the same assay format was used to determine enzyme rates at varying concentrations of ATP (2000 µM, 1000 µM, 500 µM, 250 µM, 125 µM, 62.5 µM, 31.3 µM, 15.6 µM). Data were plotted and analyzed using nonlinear regression in GraphPad Prism to extract kinetic parameters. All assays were conducted with three independent biological replicates, and error bars represent the standard deviation across these replicates.

### FITC-casein degradation assay

FITC-labeled casein (Sigma) was incubated at 37 ^*°*^C for 10 minutes in reaction buffer containing 100 mM KCl, 50 mM Tris pH 8.0, and 10 mM MgCl_2_, along with 2.5 mM ATP. Proteolytic activity was initiated by the addition of enzyme at the following final concentrations: 42 nM for wild-type LONP1, H391A/L395A, D449A, and D449A/H451A; 166 nM for H391W; and 333 nM for H391E and S453D. The release of FITC from the substrate was detected as an increase in fluorescence (excitation at 485 nm, emission at 535 nm) using a TECAN plate reader. Reactions for each variant were performed in biological triplicate. Fluorescence data were analyzed by linear regression, and the slope of each curve was used to calculate the rate of substrate degradation. Rates were normalized to wild-type LONP1 activity to enable comparison across constructs. All experiments were conducted in triplicate, and error bars represent the standard deviation of the replicates.

### Gel-based proteolytic degradation assay

Proteolysis of TFAM and StAR was assessed using gel-based assays, with substrate degradation monitored over time by SDS-PAGE. Reactions were set up by incubating 165 nM LONP1 with 5 µM of either TFAM or StAR in a buffer composed of 100 mM KCl, 50 mM Tris pH 8.0, and 10 mM MgCl_2_. This preincubation was carried out at 37 ^*°*^C for 10 minutes prior to initiating proteolysis through the addition of ATP to a final concentration of 5 mM, bringing the total reaction volume to 100 µl. At designated timepoints (0, 2, 5, 10, and 15 minutes), 10 µl aliquots were withdrawn and immediately mixed with 20 µl of Laemmli sample buffer, followed by heating at 95 ^*°*^C for 3 minutes. Samples were briefly centrifuged and 20 µl of each was loaded onto a 12% SDS-PAGE gel, which was run at 160 V for 1 hour. Protein bands were visualized by Coomassie staining using a standard solution containing 50% methanol, 10% acetic acid, and 0.25% Brilliant Blue R250.

### Sample preparation for Cryo-EM

Samples of LONP1 actively degrading substrate were made by diluting LONP1 to 30 µM with or without 5 µM of StAR, TFAM, or β-casein (Sigma) before incubating at 37 ^*°*^C for 10 minutes. Reactions were initiated by the addition of 2 mM ATP and immediately applied to grids and plunge frozen (∼ 30 seconds from reaction initiation to plunging) using a Vitrobot Mark IV system (blot time 5 seconds, blot force 1, 100% humidity, room temperature, Whatman No. 1 filter paper). All samples were applied to 300 mesh R1.2/1.3 UltrAuFoil Holey Gold Films (Quantifoil) that were glow discharged for 25 seconds at 15 mA with a Pelco Easiglow 91000 (Ted Pella, Inc.) in ambient vacuum.

### Cryo-EM data acquisition

Data acquisition was carried out on a ThermoFisher Talos Arctica transmission electron microscope operating at an accelerating voltage of 200 keV and equipped with a Gatan K2 Summit direct electron detector. Parallel beam illumination was established prior to imaging. Exposures were collected in counting mode, each lasting approximately 10 seconds at an electron dose rate of ∼7 e^-^/pixel/sec, and subdivided into ∼50 individual frames, yielding a total accumulated dose of 50 e-/Å^2^. Automated data collection was performed using the Leginon software suite,^47^ with images captured at a nominal magnification of 36,000x, corresponding to a calibrated pixel size of 1.15 Åat the specimen plane. The defocus range was set between 0.9 µm and 1.6 µm. Stage shifts were used to position the microscope over the center of 16 grid holes, followed by high-magnification image acquisition via coma-compensated image-beam-shift targeting the center of each hole. A total of 1,964 micrographs were obtained. During collection, movie frames were aligned using MotionCor2, and contrast transfer function (CTF) parameters were determined using CTFFind4^48^ within the Appion^49^ processing environment to assess image quality in real time. Additional acquisition parameters are provided in **Table 1**.

**Table 1.**
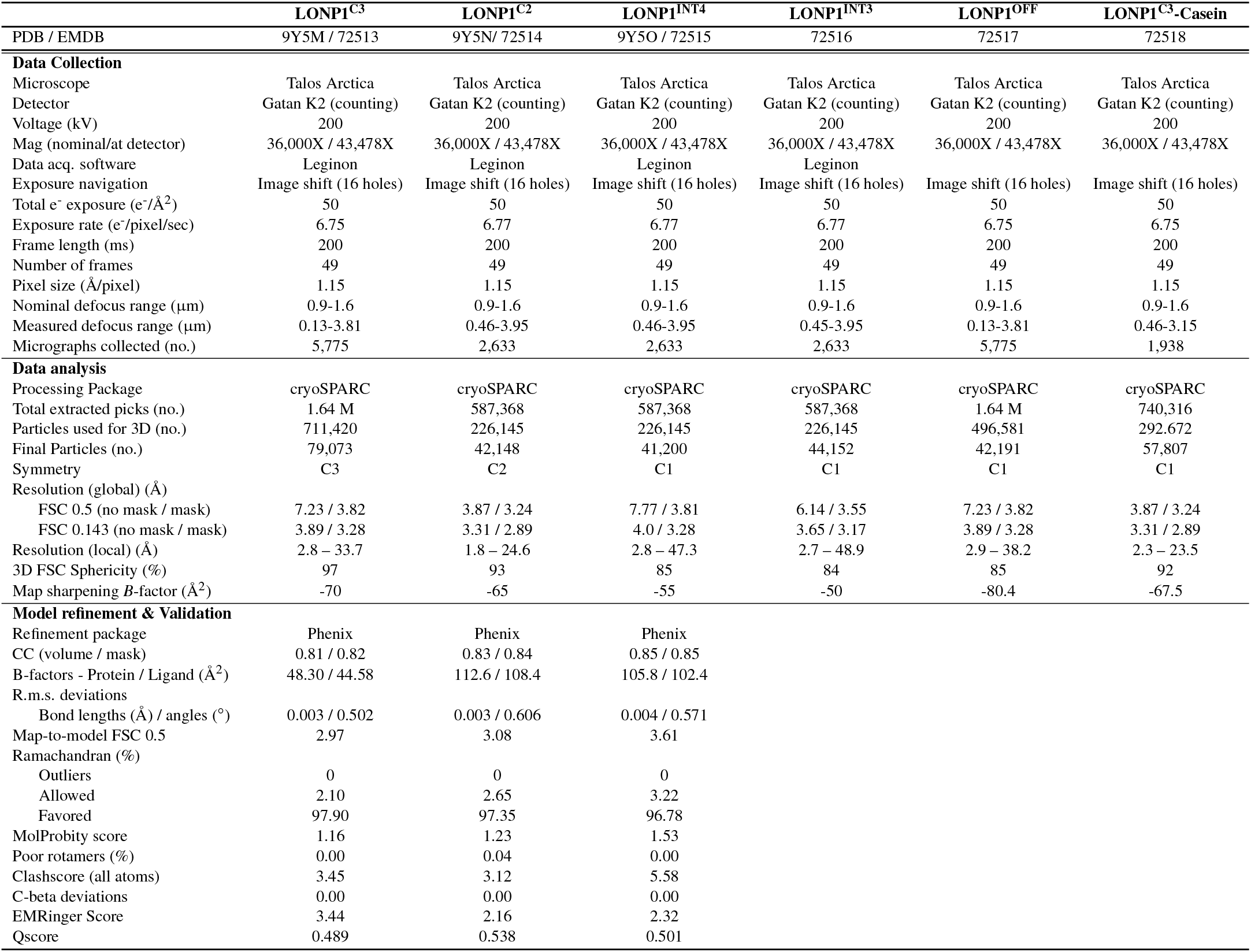
Cryo-EM data collection, image analysis, refinement, and statistics.

### Cryo-EM image analysis

The general processing strategy for all datasets is described here. Details are provided in the processing workflows in **Figs. S1, S3, S6, and S7**. Movie frames were motion-corrected and dose-weighted using MotionCor2,^50^ and the summed and dose weighted micrographs were imported into cryoSPARC v2.1-v3.3.2^51^ for patch CTF correction. Micrographs with defocus values greater than 3.3 µm or with estimated resolutions worse than 7 Åresolution were discarded. Initial picking was performed using a blob picker with a lower radius of 125 and an upper radius of 225. Particles were extracted in a 320-pixel size box without Fourier cropping. 2D classification was used to generate templates that were then used for template picking with default settings and a particle diameter of 175 Å. Template-picked particles were extracted and only the obvious non-particles consistent with ice contamination or false picks were removed using 2D classification. The initial open and closed volumes were generated using the ab initio job type with default settings and three output volumes. These were then used in subsequent heterogeneous refinement jobs to separate out the open and closed conformations and remove bad picks or damaged particles. The closed-form particle classes were then used for 3D classification using PCA initialization with default parameters except for an O-EM learning rate initialization of 0.2 and two O-EM epochs. Classes were then refined using NU-refinement refinement and the resulting models were evaluated and selecting using their resulting resolution, determined by an FSC of 0.143, and by visual interpretation of the refined maps. Final refinements were performed using NU-refinement with per particle defocus and CTF refinements enabled with standard settings. Reported resolutions were determined using the 3DFSC.^52^ Further details for the processing of each dataset can be found in the supplementary information.

### Atomic model building and refinement

Structural models of the closed conformation of LONP1 were constructed by initially fitting the previously solved LONP1^BTZ^ structure (PDB ID: 7KRZ) into the cryo-EM density maps using ChimeraX.^53^ This was followed by manually rigid body fitting specific subdomains into relevant density. Model refinement proceeded through multiple cycles of manual adjustment in Coot (v0.9.4.1EL)^54^ and automated realspace refinement in PHENIX (v1.19.2)^55^, iterating until satisfactory model-to-map correlation was achieved. To further improve geometry, minimize steric clashes, and correct unfavorable rotamers, final model relaxation was carried out using ISOLDE.^56^ Both Chimera^57^ and ChimeraX^53^ were employed for visualization, map interpretation, and figure preparation.

## Acknowledgements

We thank Will Lessin at the Scripps Research Electron Microscopy Facility for microscopy support. We thank Charles Bowman and J.C. Ducom at Scripps Research High Performance Computing core for computational support. This work is supported by the National Institutes of Health (NIH) NS095892 G.C.L. and NIH F32GM145143 to J.T.M.

Data collection used equipment supported by NIH grant S10OD032467.

## Data availability

Cryo-EM maps and associated atomic models were deposited to the Electron Microscopy Databank (EMDB) and the Protein Databank (PDB), respectively, with the following EMDB and PDB IDs: LONP1^C3^ - EMD-72513, 9Y5M; LONP1^C2^ - EMD-72514, 9Y5N; LONP1^INT4^ - EMD-72515, 9Y5O; LONP1^INT3^ - EMD-72516 (EMDB only); LONP1^OFF^ - EMD-72517 (EMDB only); LONP1^C3^-Casein - EMD-72518 (EMDB only).

## Supplementary Figures

**Figure S1.**
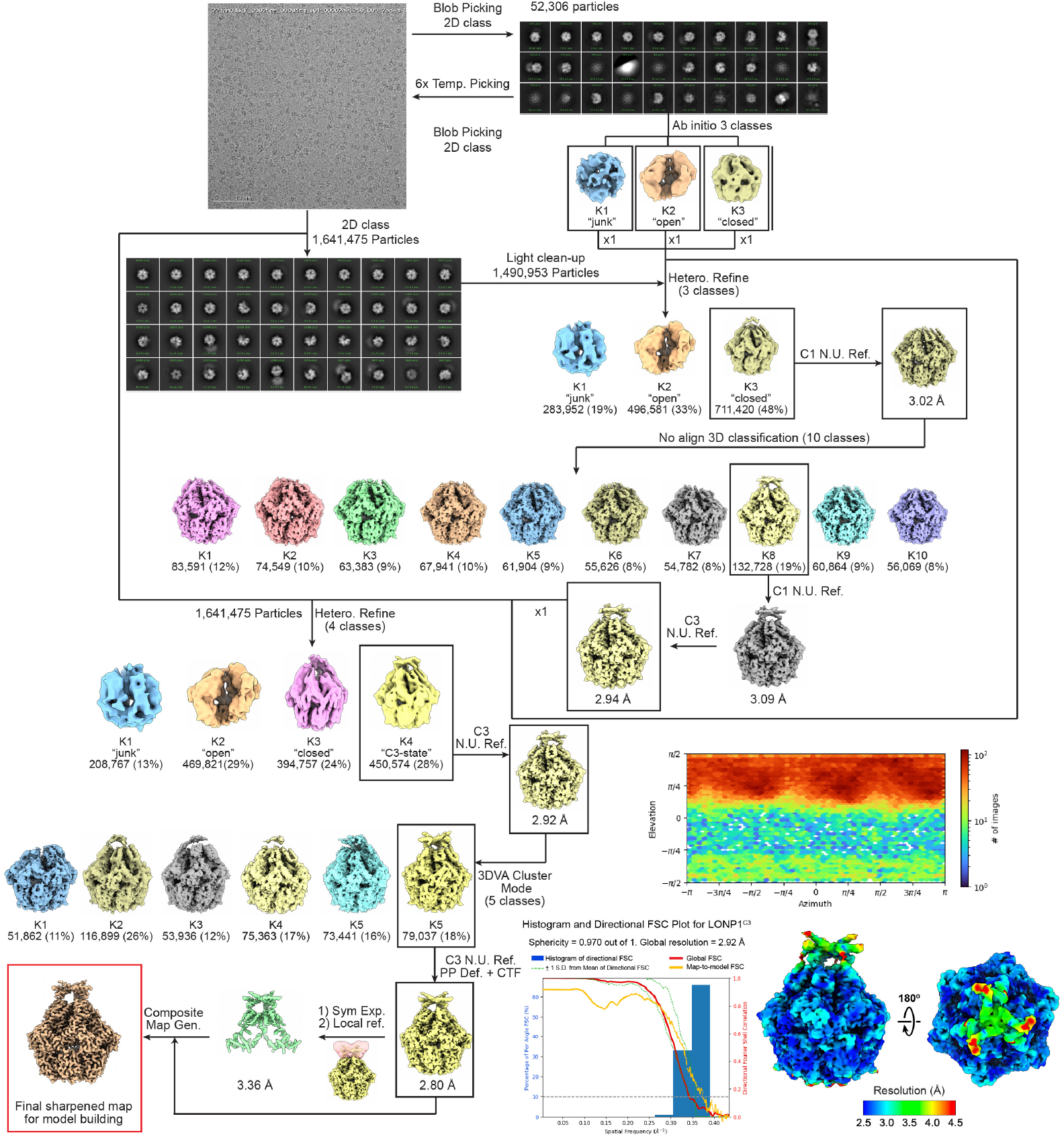
LONP1-StAR-ATP processing workflow. Representative motion-corrected, dose-weighted cryo-EM micrograph, 2D averages, and image analysis pipeline used to determine final reconstructions. In the lower right the viewing distribution plot is shown, and underneath the global FSC and map-to-model FSC is overlaid on the directional resolution histogram from the 3DFSC server^52^, alongside the local resolution map of the final reconstruction, calculated using cryoSPARC.

**Figure S2.**
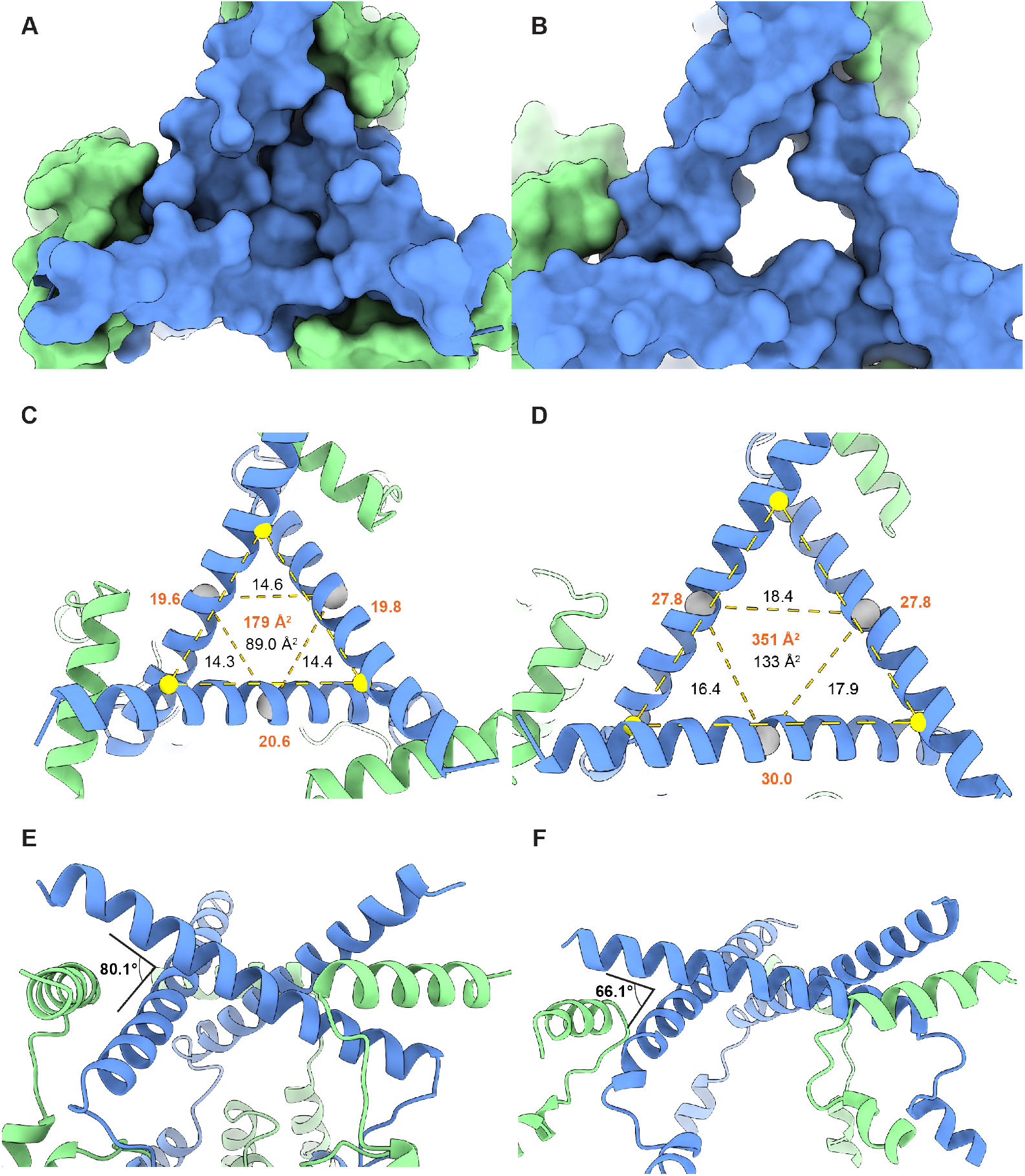
Comparison of CCD in LONP1^C3^ and substrate-bound LONP1. **A**) and **B**) Show top-view surface-representations comparing LONP1^C3^ and LONP1^ENZ^, respectively. **C**) and **D**) compare the approximate CCD dimensions of LONP1^C3^ and LONP1^ENZ^, respectively. **E**) and **F**) compare the angular pitch between CCD helices in LONP1^C3^ and LONP1^ENZ^, respectively.

**Figure S3.**
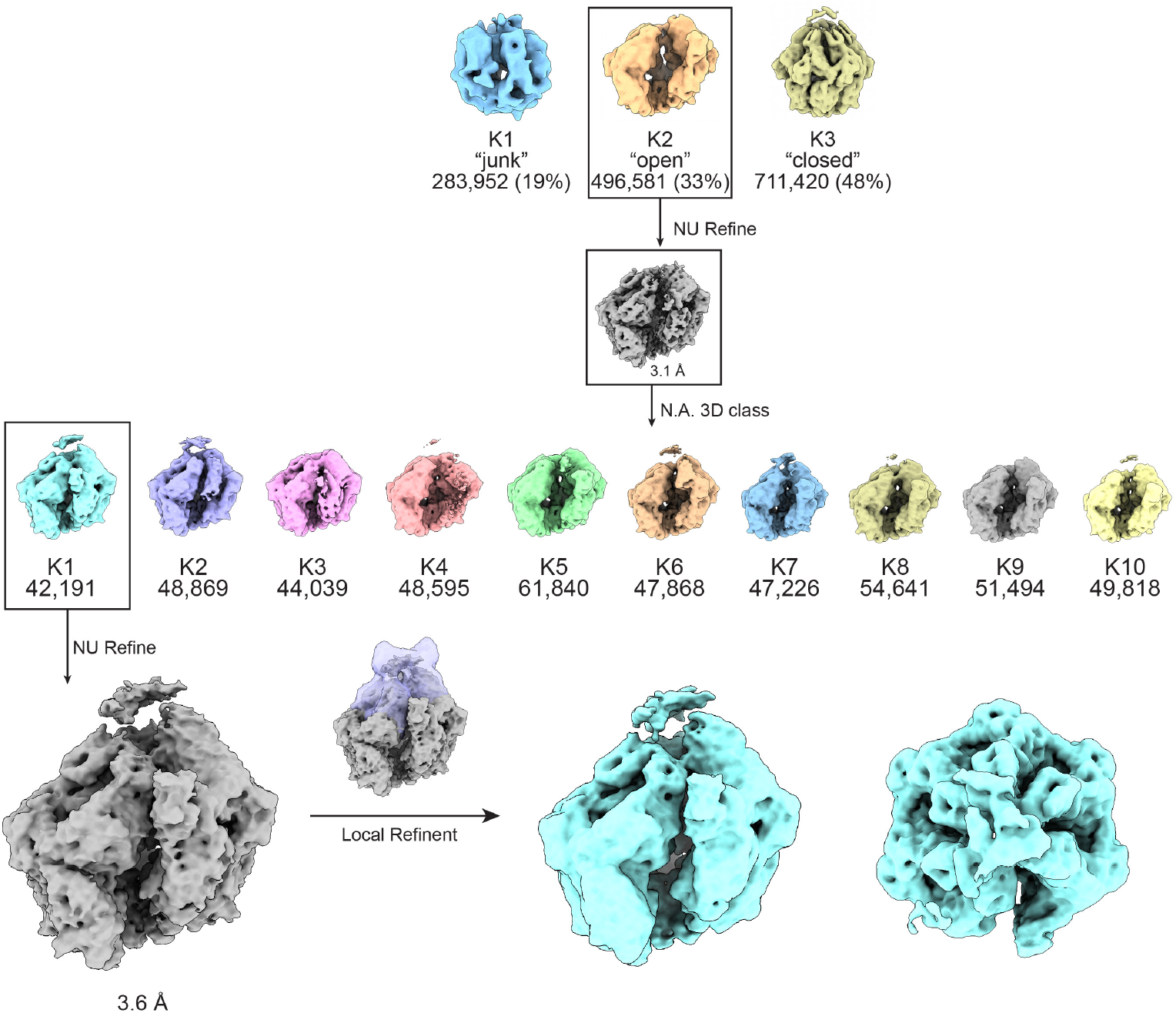
Processing workflow for LONP1OFF from LONP1-StAR-ATP dataset. This processing workflow was continued using the K2 “open” particles from the first heterogeneous refinement step shown in **Fig. S1**.

**Figure S4.**
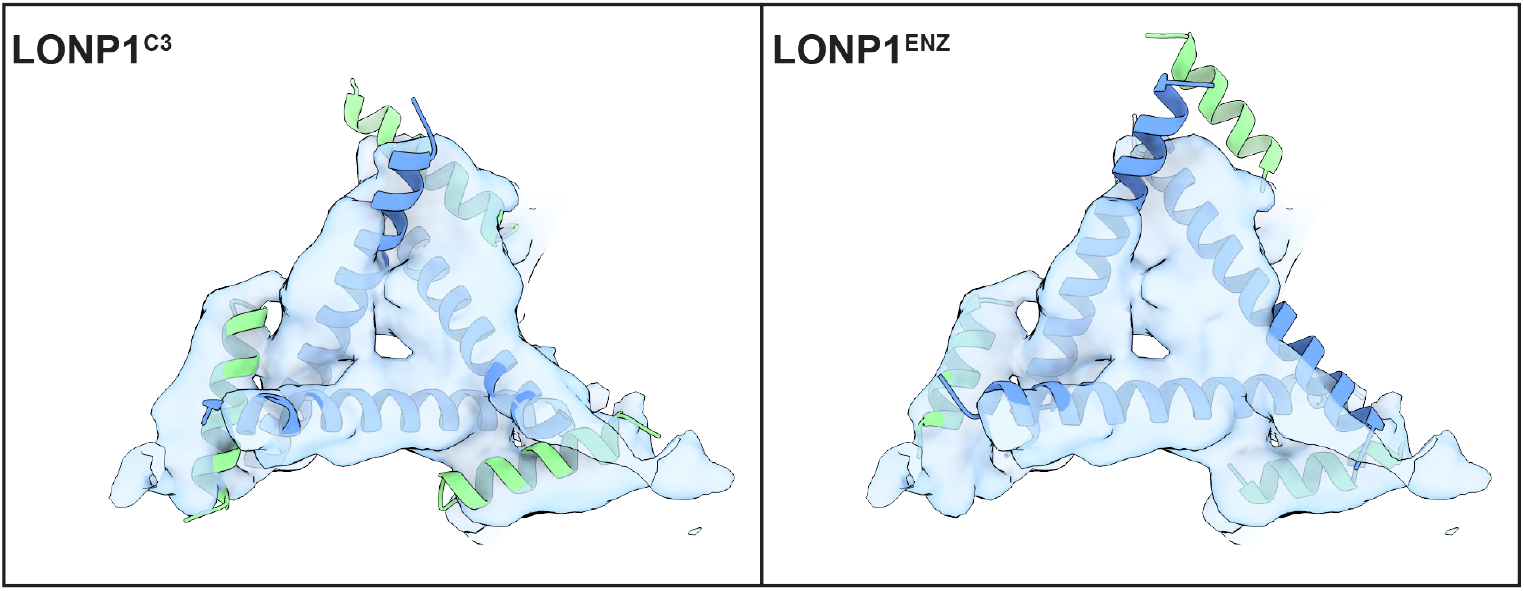
LONP1^C3^ and LONP1^ENZ^ docked into LONP1^OFF^ CCD density. The atomic models of the CCD from the LONP1^C3^ and LONP1^ENZ^ structures were rigid-body fit into the LONP1^OFF^ CCD density, showing a greater similarity to LONP1^C3^.

**Figure S5.**
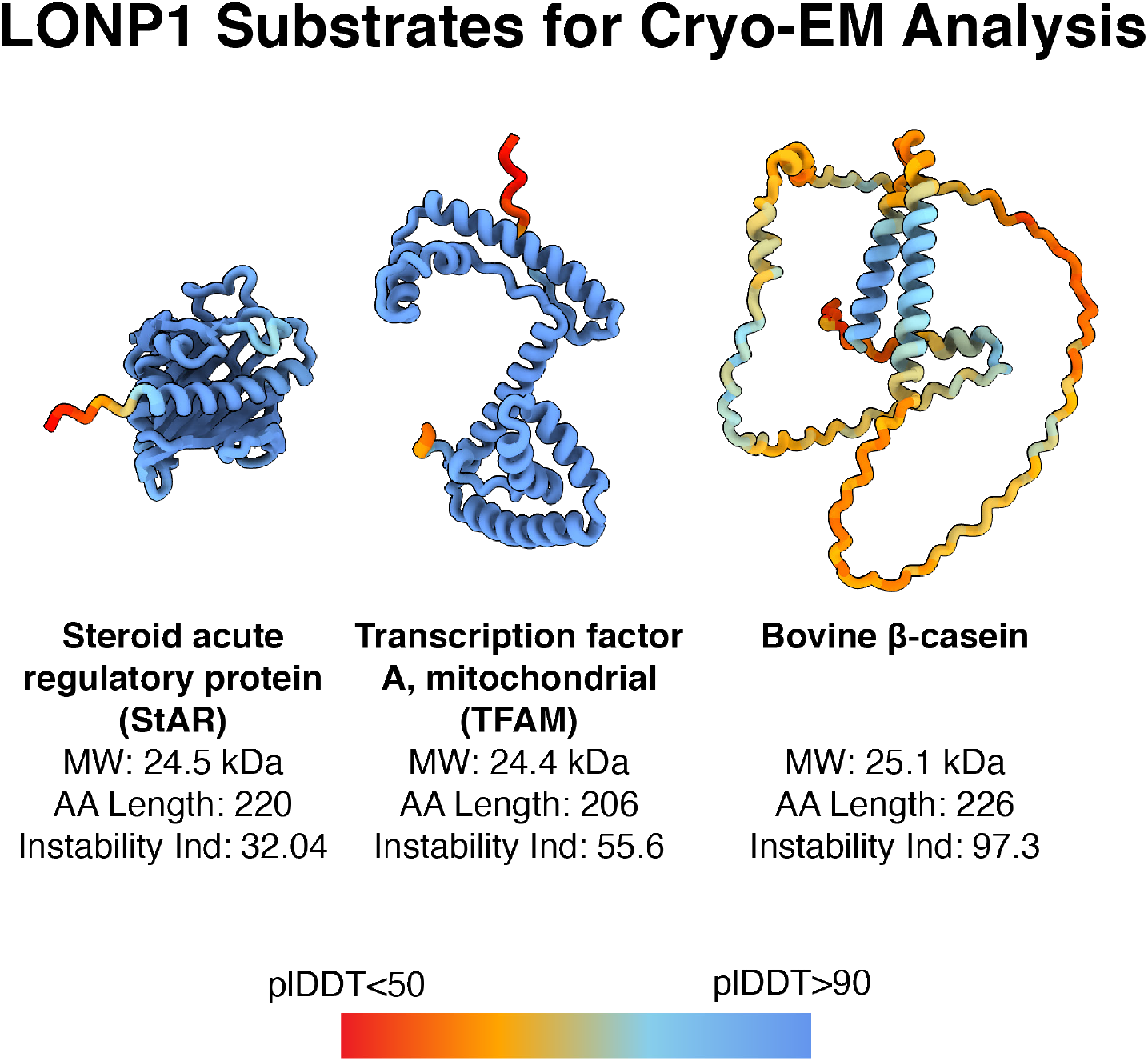
Substrate structure and stability predictions. AlphaFold2 was used to generate three-dimensional coordinates of StAR, TFAM and β-casein. The plDDT scores are mapped on top of each structure to demonstrate confidence in the predicted model. The instability index is also provided for each protein, which was calculated using ExPASY Prot Param tool.

**Figure S6.**
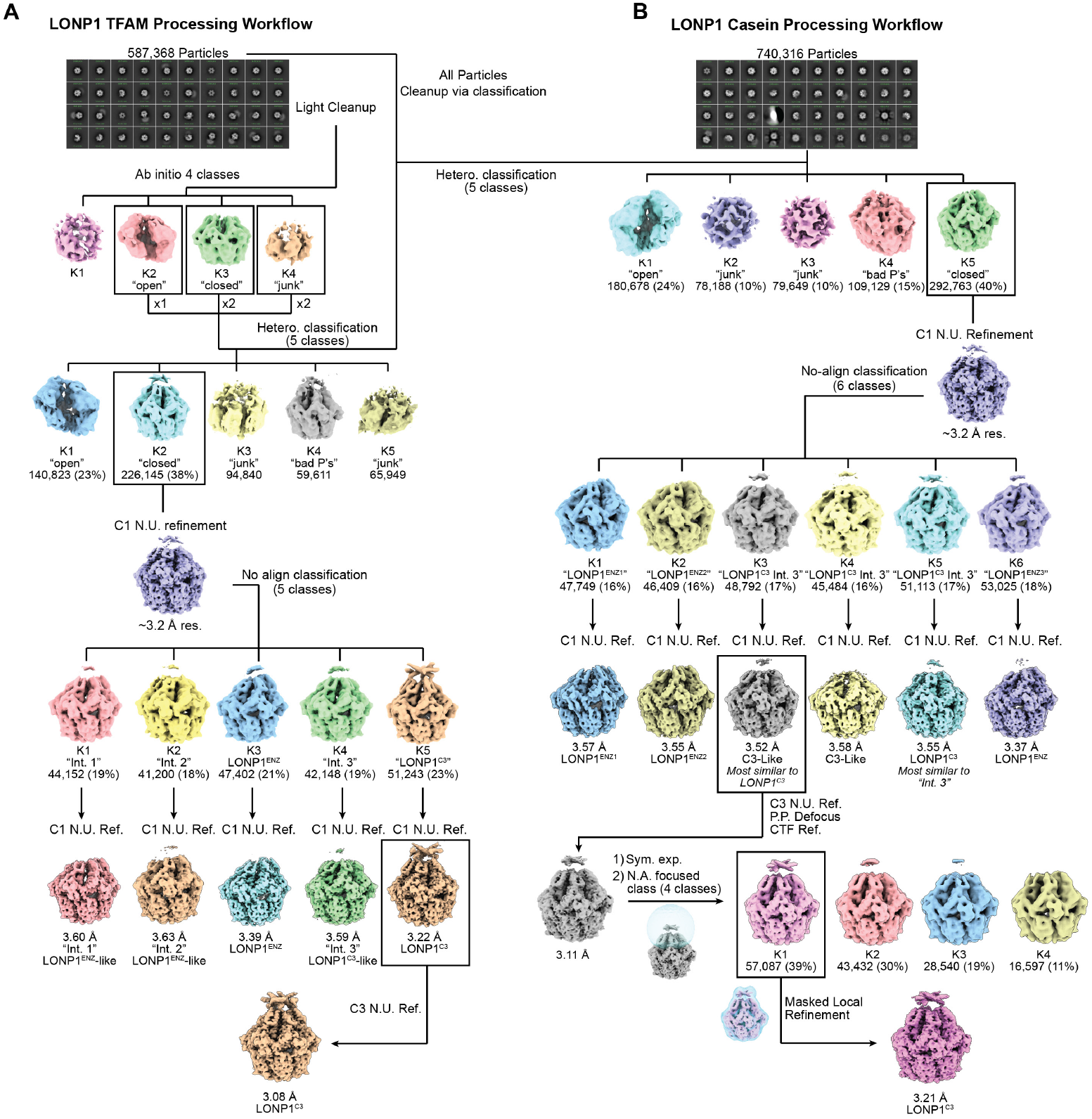
Processing workflow for A) LONP1+TFAM+ATP and B) LONP1+Casein+ATP.

**Figure S7.**
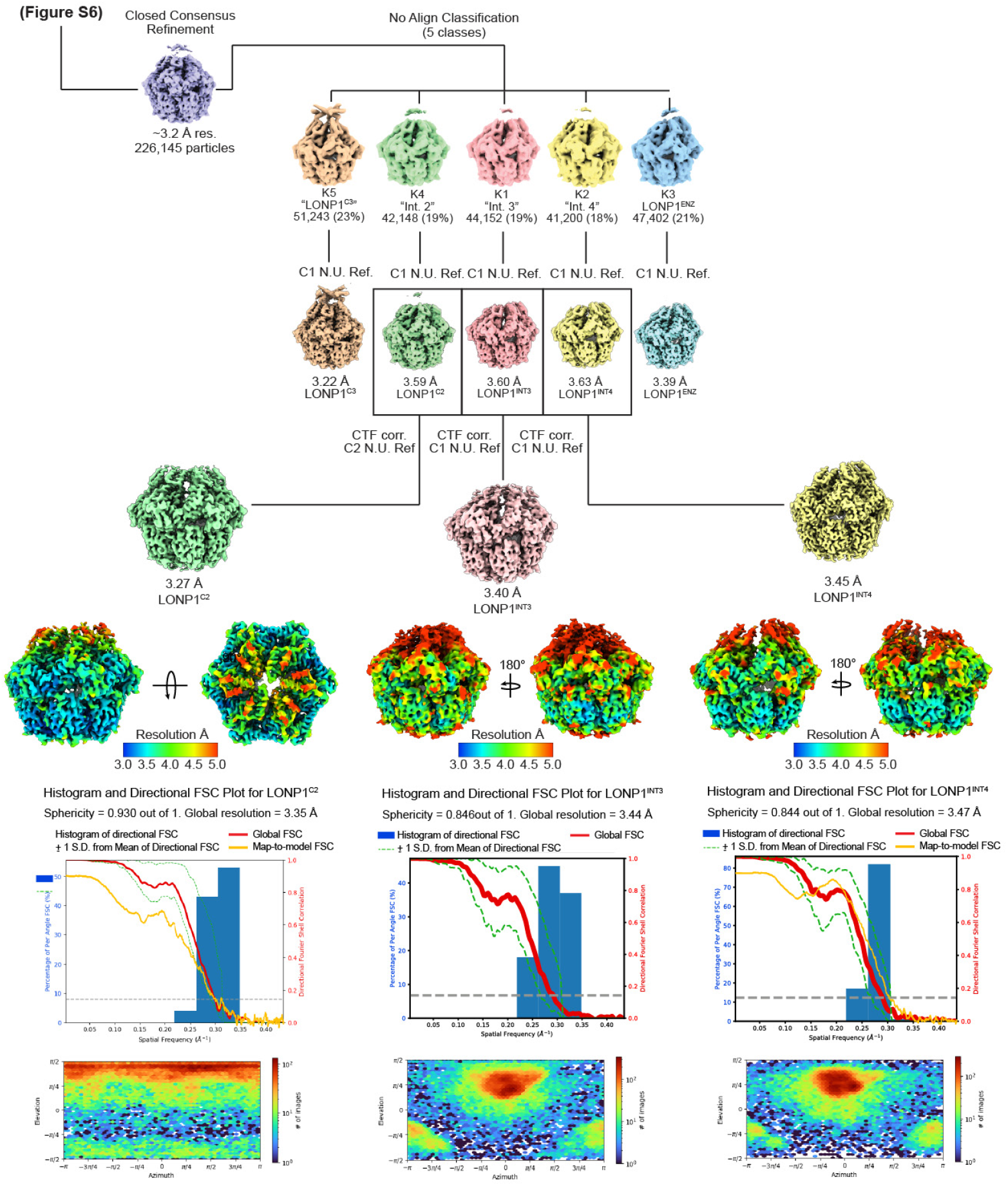
Processing workflow for LONP1^C2^, LONP1^INT3^, LONP1^INT4^. Particles from the C1 N. U. Refinement of the LONP1+TFAM workflow in **Fig. S6** were further processed to identify three intermediate conformations. For each reconstruction, the local resolution calculated using cryoSPARC is shown, as well as the global FSC and map-to-model FSC overlaid on the directional resolution histogram from the 3DFSC server. The corresponding viewing distribution plot is shown below.

**Figure S8.**
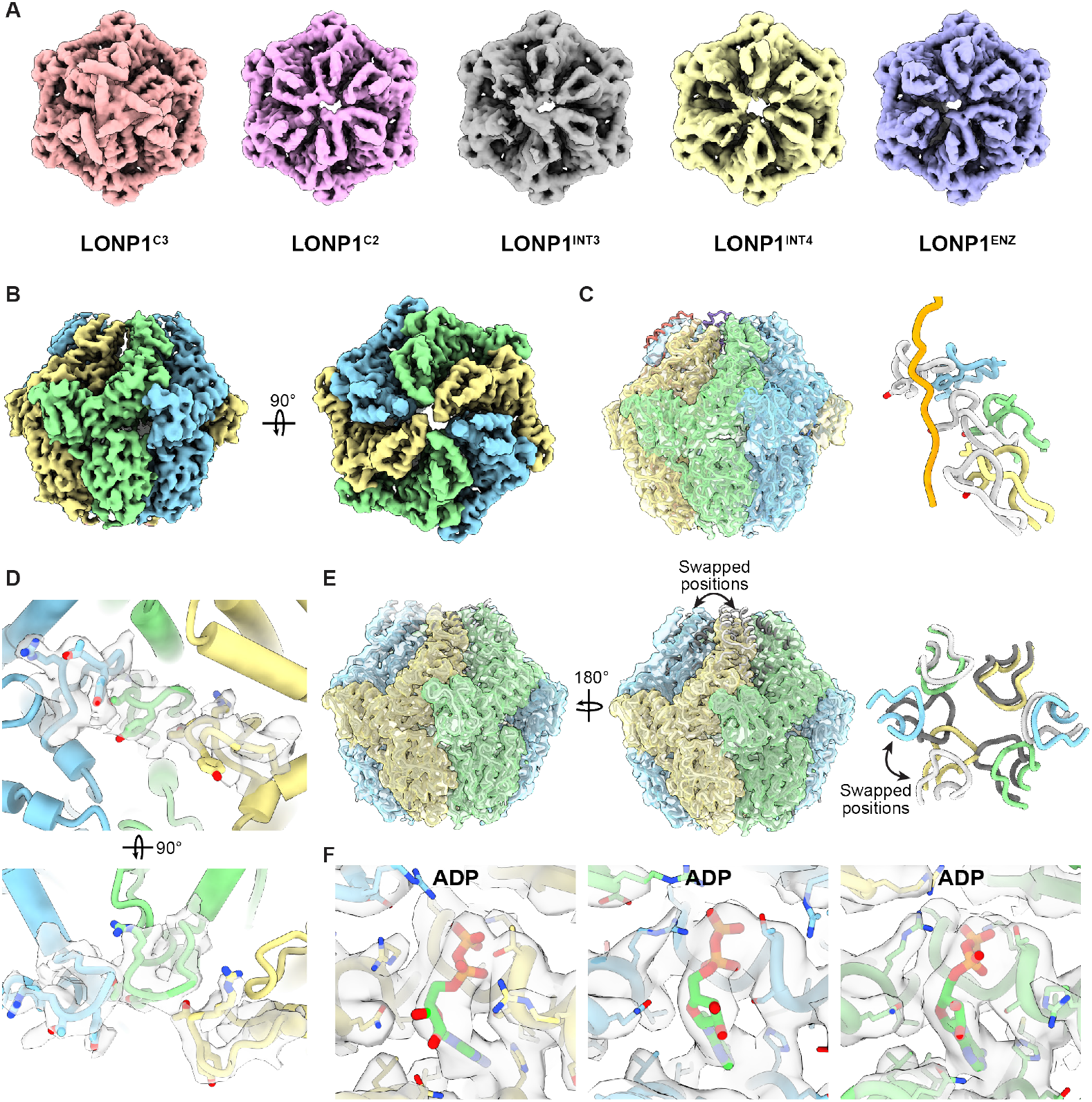
LONP1^C2^ structural analysis. **A**) Five intermediates placed in expected transition order from LONP1^C3^ to LONP1^ENZ^. LONP1^C2^ is proposed to represent the initial breaking of C3 symmetry and formation of symmetric half-spirals. **B**) LONP1^C2^ reconstruction colored by subunit position from lowest to highest (yellow, green, blue). **C**) LONP1^ENZ^ docked into LONP1^C2^ density to demonstrate that the three bottom subunits of LONP1^ENZ^ approximate the spiral forming in LONP1^C2^. An overlay of the pore loops of these positions is provided on the right hand side where the pore loops from LONP1^ENZ^ are rendered in light grey and the pore loops of LONP1^C2^ follow their conventional coloring scheme from lowest to highest (yellow, green, blue). The orange substrate is derived from the LONP1^ENZ^ model for reference. **D**) Pore loop density for LONP1^C2^. **E**) LONP1^C3^ coordinates docked into LONP1^C2^ to demonstrate that these conformations are somewhat similar and that four out of six subunits in LONP1^C2^ also have an alternating up and down arrangement. A cartoon overlay of the pore loops of LONP1^C2^ and LONP1^C3^ is provided for reference. The up and down subunits of LONP1^C3^ are colored in light and dark grey, respectively. LONP1^C2^ follows its canonical color convention denoting subunits from lowest to highest position in the spiral (yellow, green, blue). The overlay demonstrates that two subunits of LONP1^C3^ swapping position (up, down to down, up) would lead to the formation of a LONP1^C2^-like arrangement. **F**) The nucleotide density of the three subunits forming the asymmetric unit of LONP1^C2^, demonstrating that like LONP1^C3^, LONP1^C2^ is completely occupied by ADP.

**Figure S9.**
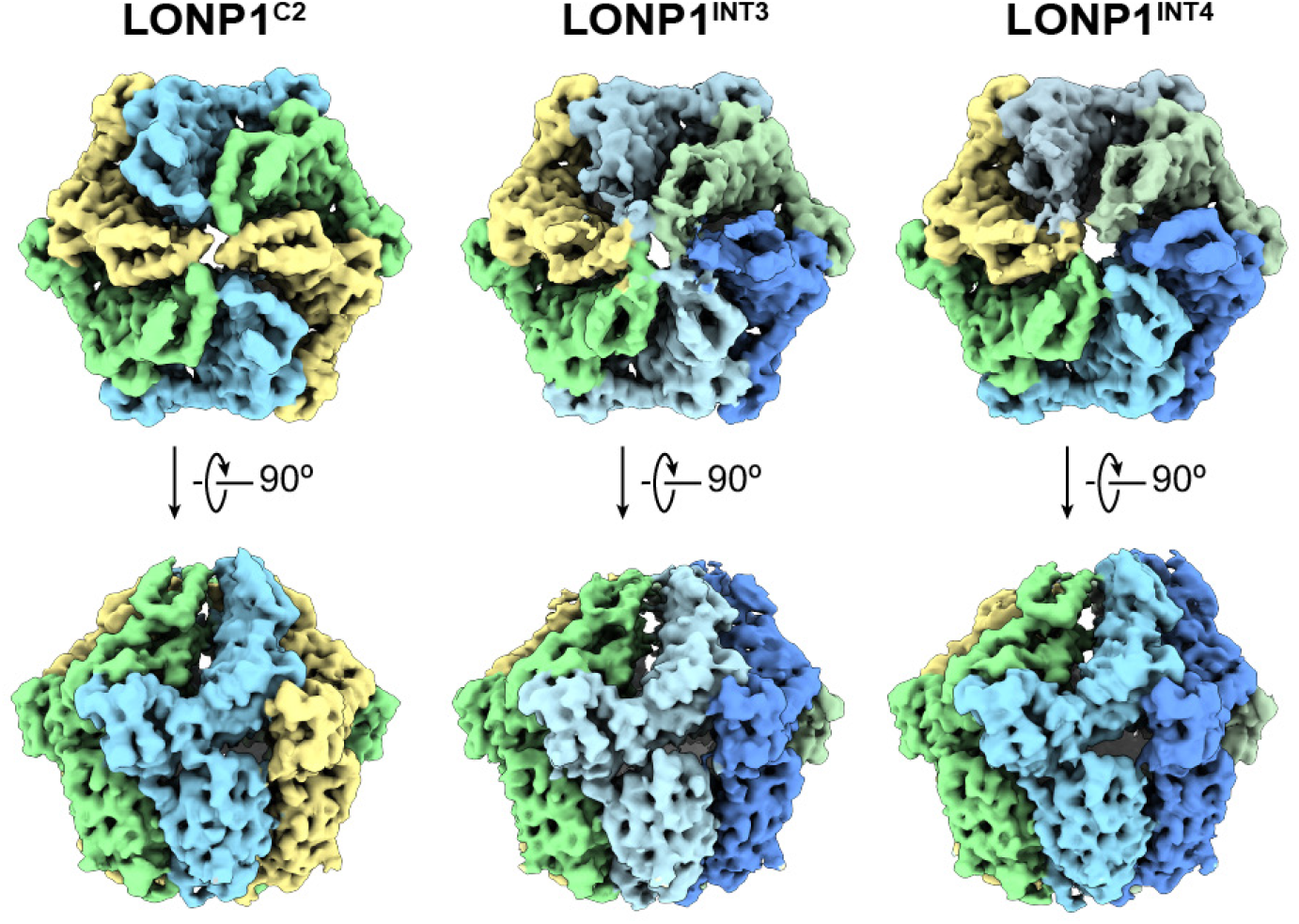
Comparison of LONP1^C2^, LONP1^INT3^, and LONP1^INT4^ reconstructions. Top and side views of LONP1^C2^, LONP1^INT3^, and LONP1^INT4^ demonstrating the progressive establishment of the fourth subunit (dark blue) in the asymmetric right-handed spiral seen in LONP1^INT3^ and LONP1^INT4^. Subunits are colored to show continuity in the progression from LONP1^C2^ to LONP1^INT4^.

**Figure S10.**
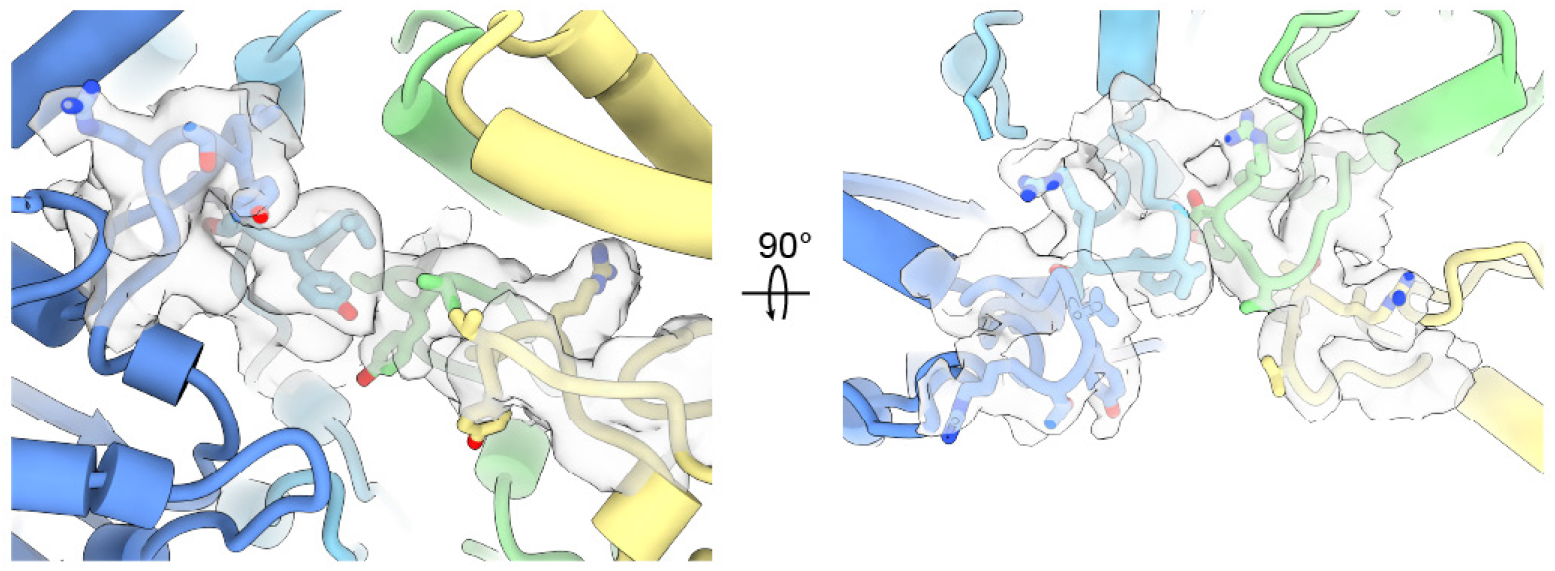
Cryo-EM density for LONP1^INT4^ pore loops. Stick representations of the pore loop residues are shown, with corresponding cryoEM density shown as a semi-transparent surface.

**Figure S11.**
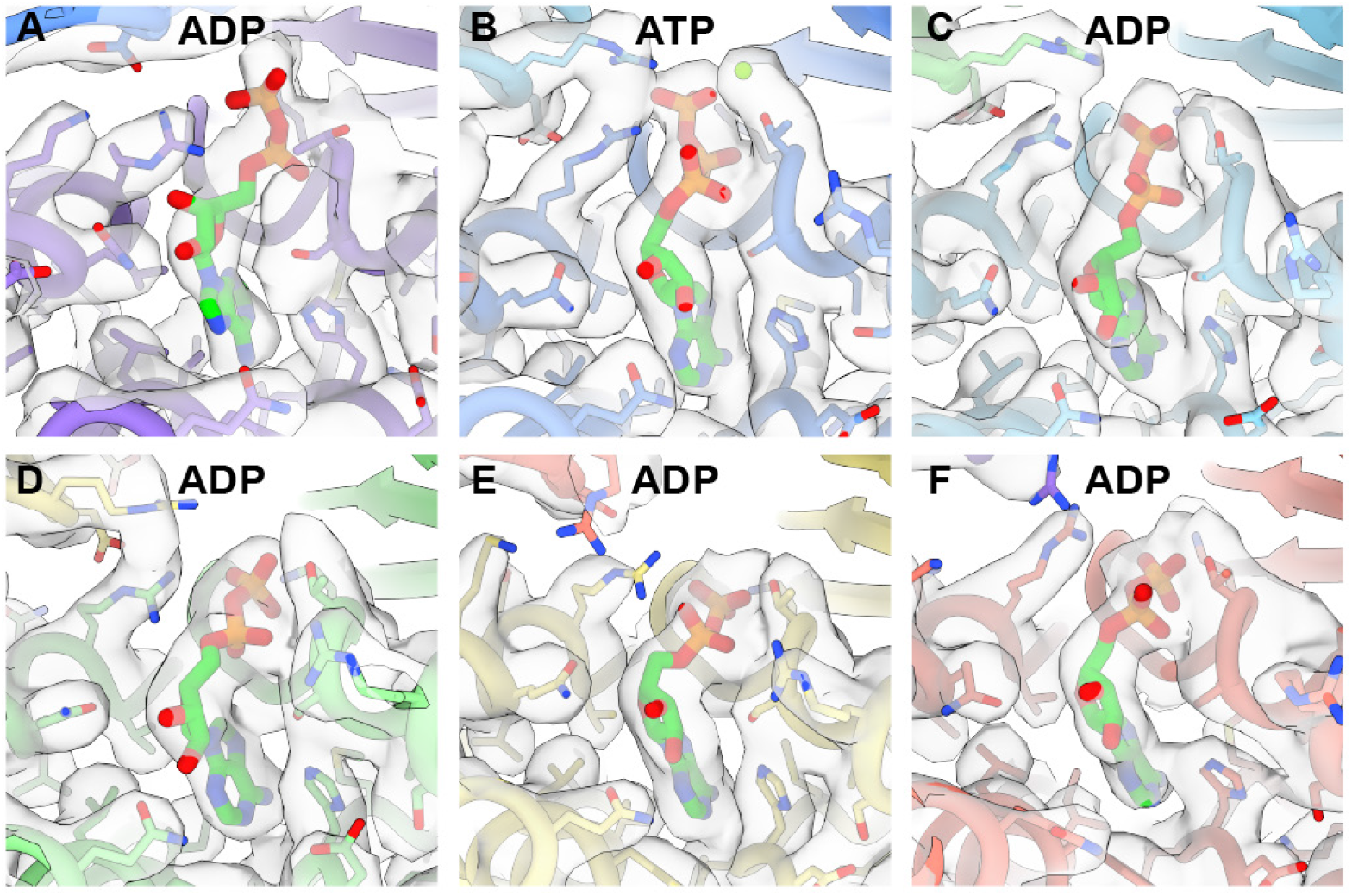
Nucleotide density for each subunit of LONP1^INT4^.

**Figure S12.**
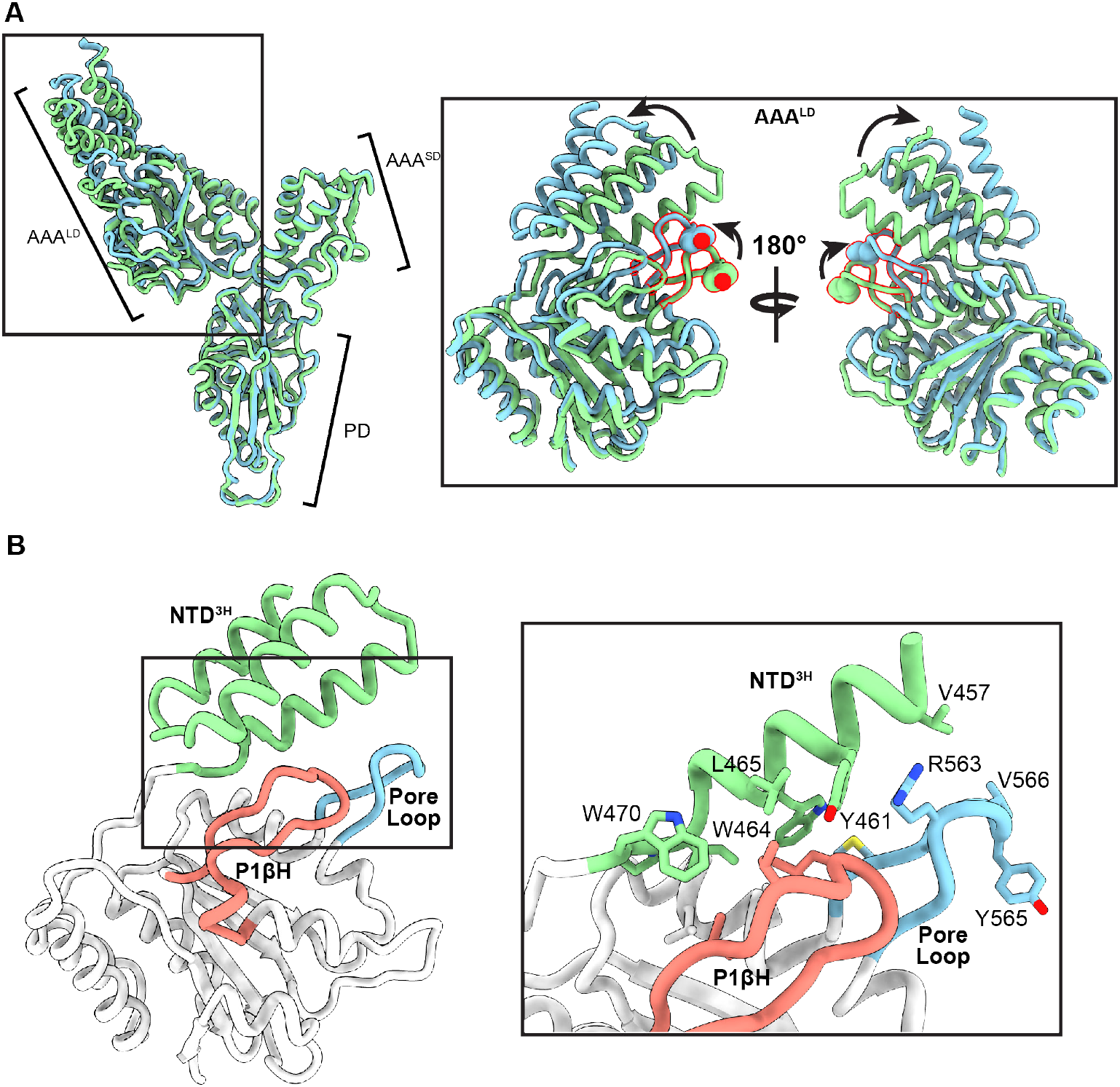
NTD^3H^ as an allosteric signal integrator. **A**) An overlay of the AAA+ domains of the up (blue) and down (green) conformation of LONP1^C3^ to demonstrate the overall architecture of the AAA+ domain and highlight regions with conformational differences between the conformers. The subdomains of LONP1 monomer are AAA+ large domain (AAA^LD^), AAA+ small domain (AAA^SD^), and protease domain (PD). The zoomed-in view provides a closer look at the AAA^LD^ demonstrating that the majority of the differences in conformation between the up and down positions in LONP1^C3^ are in the NTD^3H^, pore loops (outlined in red), and structurally associated regions. **B**) A more focused view of the AAA^LD^ highlighting key structural elements (NTD^3H^, pore loop, and presensor 1 β-hairpin (P1βH). A zoomed in view demonstrates the hydrophobic interface connecting these three regions into a unit. This network would allow conformational changes in the NTD during substrate recruitment to be communicated to key structural elements of the AAA+ via the NTD^3H^ subdomain.

